# CloneSeq: A Highly Sensitive Single-cell Analysis Platform for Comprehensive Characterization of Cells from 3D Culture

**DOI:** 10.1101/2020.11.24.395541

**Authors:** Danny Bavli, Xue Sun, Chen Kozulin, Dena Ennis, Alex Motzik, Alva Biran, Shlomi Brielle, Adi Alajem, Eran Meshorer, Amnon Buxboim, Oren Ram

**Affiliations:** Department of Biological Chemistry, Alexander Silberman Institute of Life Sciences, The Hebrew University, Jerusalem, Israel; Department of Genetics, Alexander Silberman Institute of Life Sciences and The Edmond and Lily Safra Center for Brain Sciences (ELSC), The Hebrew University, Jerusalem, Israel; Alexander Grass Center for Bioengineering, Benin School of Computer Science and Engineering, Hebrew University of Jerusalem, Jerusalem, Israel; Department of Cell and Developmental Biology, Hebrew University of Jerusalem, Givat Ram, Jerusalem, Israel

**Keywords:** CloneSeq technology, Cancer heterogeneity, 3D culturing, Cancer clonal expansion, Cellular stemness, Clone-to-clone variation, Single-cell RNA-seq

## Abstract

Single-cell assays have revealed the scope and importance of heterogeneity in many biological systems. However, in many cases, single cell limited sensitivity is a major hurdle for uncovering the full range of cellular variation. To overcome this limitation, we developed a complementary single cell technology, CloneSeq that combines clonal expansion under controlled culture conditions inside three-dimensional (3D) hydrogel spheres and droplet-based RNA sequencing (RNA-seq). We show that unlike single cell transcriptomes, clonal cells maintain cell states and share similar transcriptional profiles. CloneSeq analysis of Non-small-cell lung carcinoma (NSCLC) cells revealed the presence of novel cancer-specific subpopulations, including cancer stem-like cells (CSLCs). Standard single cell RNA-seq assays as well as cell-to-clone tracing by genetic barcoding failed to identify these rare CSLCs. In addition to CSLCs, clonal expansion within 3D soft microenvironments supported cellular stemness of embryonic stem cells (ESCs) that retained their pluripotent state in the absence of pluripotent media and improved epigenetic reprogramming efficiency of mouse embryonic fibroblasts. Our results demonstrate the capacity of CloneSeq, which can be effectively adapted to different biological systems, to discover rare and previously hidden subpopulations of cells, including CSLCs, by leveraging the broader expression space within clones.

## Introduction

Single-cell studies have revealed that there is considerable cell-to-cell variation within tumors of different cancer types^1–4^ and during embryonic stem cell (ESC) differentiation^5,6^. For example, single cells derived from glioblastomas have inherent variation in their transcriptional expression^7^, and lung adenocarcinoma cells have heterogeneity of immune response-related gene expression^8^. Other cancer-related single-cell studies have characterized infiltrating immune cells such as T cells, which have furthered our understanding of their heterogeneous organization, clonal expansion, migration, and functional-state transitions^9^. As cells from cancerous tumors have an assortment of cellular mutations that give rise to cells with the ability to de-differentiate, tumors are highly heterogeneous, making single cell-based profiling a powerful tool to dissect the underlying cellular structures. Similarly, investigations of early development and differentiation of stem cells greatly benefit from single-cell resolution approaches. ESCs grown *in vitro* perpetuate the broad developmental potential of naive founder cells in the preimplantation embryo^10^. ESCs are composed of cells in different states and differentiation potentials^11^; however, limitations in the sensitivity of single-cell technologies hinder our ability to understand the fine-tuning of cellular hierarchies. Technological advances such as high-resolution cell imaging^12^ and single-cell profiling of epigenomic and genomic sequences^13,14^ suggest that cellular heterogeneity results from more than just a mixture of different cell types. It appears that a given cell type can be composed of a subtle assortment of cells with different states. These different states allow cells to adjust to changing conditions by committing to a certain differentiation trajectory or simply minimizing metabolism. This, in turn, appears to enhance drug resistance development, as recently suggested for acute myeloid leukemia cells^15^.

Stochasticity, or randomness, is a strong component in the accumulation of cellular variation^16^. Stochastic effects are very difficult to study as in many cases they describe chaotic processes we do not understand and that are difficult to separate from biological and technical noise. Stochastic effects and noise often confound single-cell measurements. For example, the nonlinearity of transcription, also known as transcriptional bursting, can lead to errors in clustering of data^17^. Cell-cycle regulation is an important biological feature, but if cells are in different stages of the cell cycle, alterations in cell-cycle regulators and cycling genes can mask subtle differences that determine distinct cellular states^18^. Uneven culturing conditions, in terms of the distribution of reagents and oxygen and differences in surface tension and elasticity, can affect cellular outcomes. Moreover, the presence of inevitable technical variability introduced during sample processing steps also causes batch effects^19^. Finally, single-cell profiling techniques inherently suffer from low sensitivity that can lead to false-negative and false-positive results^20^. These confounders can strongly influence the level of randomness attributed to measurements, thus single-cell experimental data are highly noisy and difficult to interpret, especially in the context of cellular states.

To overcome these hurdles, we developed CloneSeq, a 3D clone-based RNA-seq approach. Our hypothesis was that clones are composed of cells more similar to each other than cells picked at random, and that analysis of clonal cells would have improved sensitivity and broader coverage relative to single-cell RNA-seq (scRNA-seq). Our results support this hypothesis, as cells originating from a given clone had more similar transcriptional profiles than cells across clones. The small clones in our 3D system also had detectably different phenotypes. We leveraged this observation to perform an in-depth dissection of cellular heterogeneity in lung adenocarcinoma PC9 cells. We were able to characterize different cellular states including cancer stem-like cells (CSCs), high and low replicative cancer cellular states, and different levels of invasiveness. Such features cannot be detected using scRNA-seq due to its low mapping resolution. As our 3D culturing method supports cancer cell growth and nourishes cellular stemness, it could be optimized for primary tumor cell expansion. Finally, we show that this 3D system induces embryonic stem cell formation without standard supplements, such as LIF and MEK and GSK3 inhibitors (2i), and improves efficiency of induced pluripotent stem cell (IPSC) production, making it superior than standard ESC culturing methods.

## Results

### Developing a 3D hydrogel cell culture system

To support the expansion of single cells into small clones in a confined, controlled, and robust setting, we developed CloneSeq, a method that supports spherical 3D tissue culturing and sequencing. The method includes three steps: 1) Capturing of single cells inside soft, biocompatible, and biodegradable hydrogel spheres using a microfluidic device; 2) clonal expansion of single cells within the hydrogel spheres up to 60 cells, depending on cell type and size; and 3) performing single-clone RNA sequencing by uniquely barcoding each clone inside nanoliter droplets. Since single cells are allowed to expand within the hydrogel spheres while maintaining inter-clonal cell states, single-clone transcriptional profiling facilitates the detection of a large number of genes (increased coverage), including genes that are lowly expressed (increased sensitivity), without averaging out transcriptomes such as in cell population RNA-seq.

We optimized a microfluidic architecture for capturing single cells within 3D hydrogel spheres (**Figure 1A**). The inflow of cells and maleimide-dextran (MALDEX) precursor is first mixed with polyethylene glycol dithiol (PEGDT) at the first junction (**Figure 1A–1**). Small nanoliter aqueous MALDEX-PEGDT droplets are formed via oil-phase “pinching”, with one cell per 5 droplets encapsulation rate (**Supplementary Table 1**) MALDEX polymerization and PEGDT crosslinking occurs spontaneously via thiol-maleimide click chemistry without reacting with the cells.^21^ Gelation is completed within ten seconds^22^ The device is designed to generate ~700 hydrogel spheres per second with a diameter of 60 ± 3 μm. The cured gels are then collected into a tube and immersed in culture medium (**Figure 1A–2**), permitting the diffusion of growth factors and supporting the proliferation of encapsulated cells.

**Figure 1.**
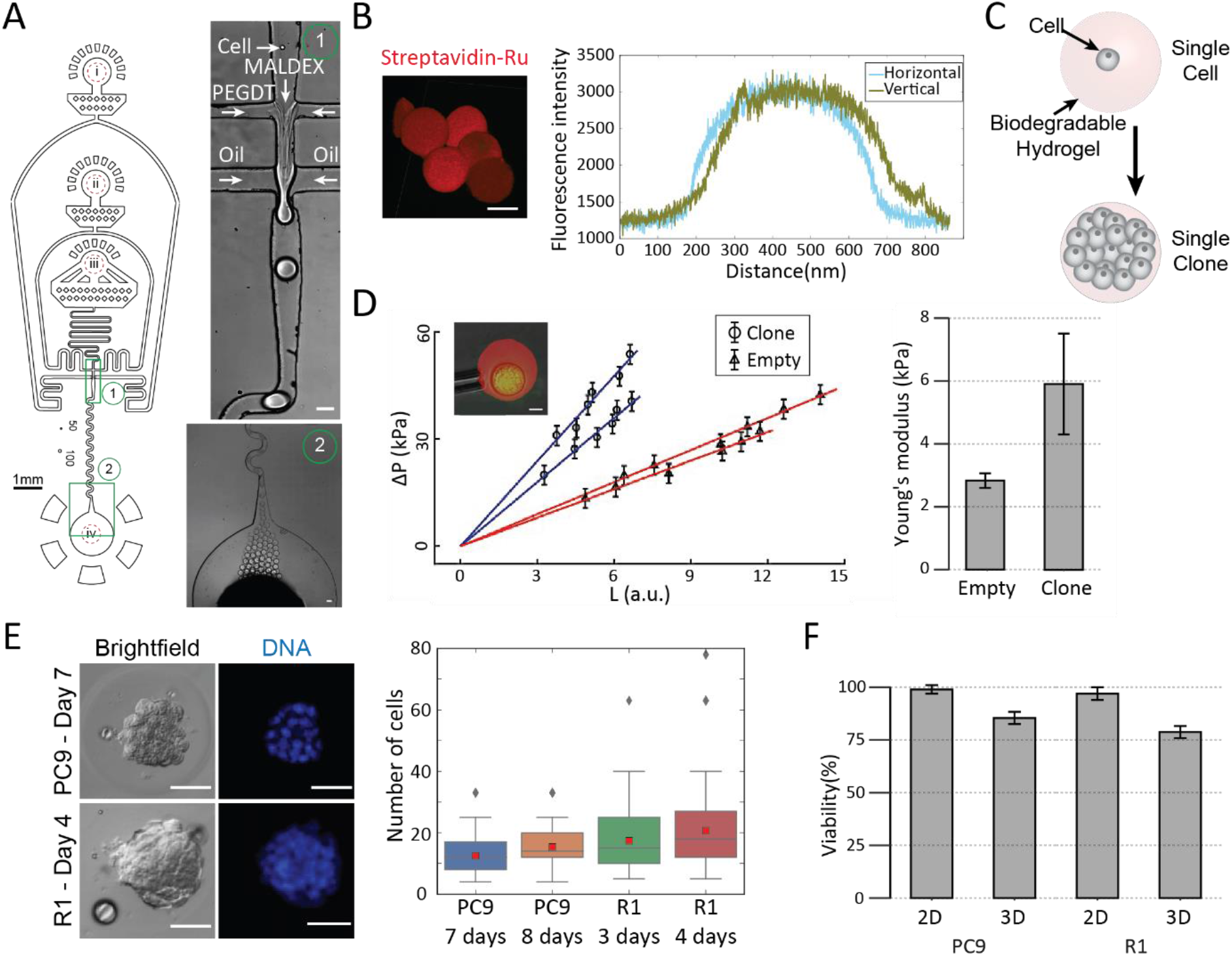
Cell encapsulation, clone formation and characterization. (**A**) Clone encapsulation. Single cells were encapsulated within biodegradable hydrogel spheres using a microfluidic device (left). The microfluidic device consists of (i) a carrier oil inlet, (ii), a PEGDT inlet (iii) a cell-MALDEX precursor mix inlet, and (iv) the droplet collection outlet. Zoom-in of (1) encapsulation and (2) outlet regions are shown on the right (scale bar: 50 μm). Microfluidic channels depth: 50 μm. (**B**) Confocal imaging (left, scale bar: 50 μm) and fluorescence cross-sectional profiles (right) of the streptavidin-RU staining show round shapes with equal density across the hydrogel spheres. (**C**) Using the cell type specific medium, the PEGDT-MALDEX hydrogels support the proliferation of encapsulated cells and clone formation. (**D**) Left: The mechanical properties of empty hydrogel spheres and clones were evaluated using micropipette aspiration. Scale bars: 10 μm. Left: the aspirated length (L) of empty hydrogel spheres (n=2) or a clone (n=2) increases linearly with applied suction pressure (ΔP) indicative of a purely elastic response. Right: Elasticities (as Young’s modulus) calculated based on pipet aspiration test (n=2). Hydrogel spheres encapsulating clones of PC9 cells are stiffer than empty hydrogel spheres. (**E**) Left: Bright-field and fluorescence microscopy images show the formation of 3D clones of encapsulated PC9 cells and R1 ESCs. Scale bars: 30 μm. Right: Number of cells per clone of PC9 and R1. (**F**) Viability comparison of PC9 and R1 ES cells cultured on gelatin-coated plates (2D) and inside 3D hydrogel spheres (3D). (n=2)

To verify structural homogeneity and uniformity, we modified the MALDEX backbone with thiolated biotin and stained the hydrogels spheres with rhodamine-conjugated streptavidin (streptavidin-RU). The high affinity of the biotin-streptavidin interaction ensures that once formed, the specific staining will not be affected by changes in pH or rinsing^23^. Indeed, confocal microscopy cross-sectional imaging revealed a uniform distribution of streptavidin-RU within the PEGDT-MALDEX hydrogel spheres (**Figure 1B**). Next, we characterized the mechanical properties of the PEGDT-MALDX hydrogel spheres using micropipette aspiration^24^. The spheres deformed elastically in response to applied stresses, reaching a steady state aspiration length that correlated linearly with the applied intra-pipette pressure (**Figure 1D**). Using the homogenous half-space model approximation^25^, we calculated the stiffness of the hydrogel spheres. The elasticity of the spheres was 3 kPa, which is consistent with the microelasticity of soft tissues such as fat and kidney^26^.

Owing to their dextran backbone, the PEGDT-MALDEX hydrogel spheres can be remodeled by the encapsulated cells. The spheres were shown to support the viability and proliferation of several cancer cell lines including PC9 absent of extracellular adhesion ligands (**Figure 1E**). PC9 clones consisted of 12 cells and 15 cells on average after 7 and 8 days in culture, respectively (**Figure 1E**), and maintained the viability of 87% of the cells within the hydrogel (**Figure 1F**). The expansion of PC9 clones increased the effective stiffness of the hydrogel spheres from 3 kPa to 6 kPa, which is comparable with lung tissue mechanics consistent with PC9 tissue origin (**Figure 1D**)^26^. The increase in the effective stiffness of the spheres is attributed to the internal prestress that is generated by the contractile cells of the encapsulated clone. Unlike PC9 cells, supporting the viability and proliferation of ESCs within the hydrogel spheres required the insertion of cell adhesion signals and cleavable properties by cell-secreted metalloproteinase (MMP)^27,28^. Hence, we encapsulated ESCs within hydrogel spheres which were supplemented with thiolated RGD peptides that mediate cell adhesion to the gel via integrin transmembrane receptors^27^. Additionally, the PEGDT crosslinker was replaced by a dithiolated PEG-peptide conjugate (MPEGDT) crosslinker, using an amino acid sequence motif (PLGLWA) that serves as a cleavable site for matrix metalloproteinase (MMP)^27,28^. Indeed, the MPEGDT-MALDEX hydrogel spheres supported ESC proliferation and the generation of ESC clones consisting of 15 cells and 20 cells on average after 3 days and 4 days in culture, respectively (**Figure 1E**). Specifically, after 4 days ESCs reached clone size of approximately 50% of the hydrogel diameter, while maintaining the viability of 77% of the cells (**Figure 1F**).

### CloneSeq: profiling of clones using modified inDrops protocol

For mRNA profiling of clones, we designed a microfluidic device to capture clones in drops and barcode their mRNAs using a modified inDrops protocol^29^ (**Figure 2A**). The device consists of two junctions: one that combines the lysis, barcodes, and clone suspension, and another junction for encapsulating the aqueous inputs in the oil phase. To allow the clones to flow smoothly, the height of the device was set to 120 μm, producing an encapsulated aqueous phase of 3.3 ± 0.2 nl. To assess clone-to-clone variation, we considered each 3D spherical hydrogel spheres containing one clone as a singular entity. To extract mRNA from the clones, hydrogel spheres were dissolved using dextranase. During the development of the method, we found that the droplet-based reverse transcription reaction used in the standard inDrops^6^ protocol was inhibited in the presence of dextranase (**Supplementary Figure 2**). However, an attempt to lyse the cells within the hydrogel spheres without dextranase, which would have allowed mRNA to diffuse out of the hydrogel spheres through pores and bind the barcodes inside drops, significantly reduced the number of RNA molecules captured (**Figure 2B**).

**Figure 2.**
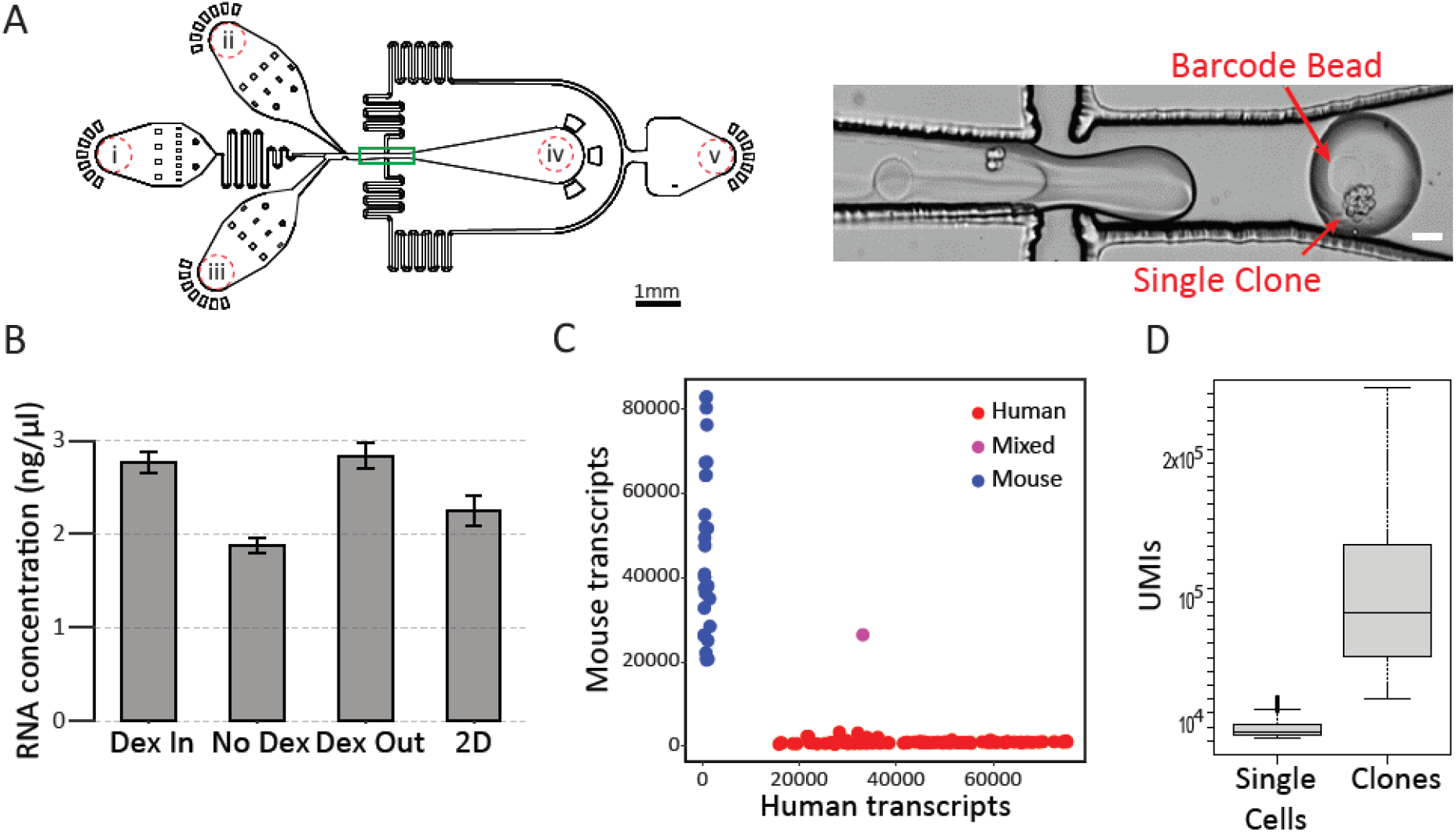
CloneSeq increases sensitivity and coverage of transcriptional profiling. (**A**) Total RNA barcoding of encapsulated clones was performed within a modified *InDrop*-based microfluidic configuration. Microfluidic channels depth: 120 μm. Zoom in of the encapsulation junction (green frame) shows a single clone co-encapsulated with a barcoded bead immersed in lysis buffer. (**B**) Optimization of mRNA extraction from PC9 cells in bulk. mRNA was extracted from cells that were embedded inside a PEGDT-MALDEX hydrogel using lysis buffer supplemented with dextranase (*Dex In*), using lysis buffer only (*No Dex*), and following a twostep approach in which dextranase was first used for degrading the gels and releasing the cells followed by lysis buffer (*Dex Out*). As a control, mRNA was also extracted from a standard 2D culture of PC9 cells (*2D*). Cell number was maintained equal for all conditions. Biological replicates: n = 2. (**C**) CloneSeq of human and mouse clones maintains organismal origin and excludes mixing of multiple clones per droplet and multiple droplets per a barcoded bead. (**D**) The number of non-redundant transcripts with unique molecular identifier (UMI’s) is threefold greater compared with standard InDrop single-cell RNA-seq.

Next, we tested the Drop-seq protocol^30^, in which cell lysis and mRNA-capture occurs inside drops and the reverse transcription (RT) reaction occurs after the drops are dissolved. Using this protocol, the hydrogel spheres are dissolved in drops by dextranase after the RT reaction is complete. However, we observed that given similar input cell types and the similar concentration of drops, Drop-seq of clones did not produce significantly higher numbers of transcripts compared to Drop-seq of single cells. We suspect that this limitation is due to the fact that each commercially available barcoded bead contains a fixed number of barcoded primers (~10^6^)^30^ (**Supplementary Figure 3**).

To overcome the RT inhibition observed with the inDrops protocol, we took advantage of the observation that when cells grow inside hydrogel spheres, they tend to form highly adhesive and stable structures and remain as spheroids even after the surrounding hydrogel spheres has been dissolved. Therefore, in order to expose the RT to dextranase, the hydrogel spheres were dissolved with dextranase before being introduced into the microfluidics apparatus (**Figure 2A**). After dextranase treatment, the clones were resuspended in 10% OptiPrep Medium in PBS and then introduced into the device. This inDrops-based “dextranase-out” protocol also allowed us to control the size of the polyacrylamide-based barcoded beads and hence to increase the number of barcoded primers by increasing the diameter of the polyacrylamide beads and/or increasing primer concentration as previously described ^30^. The number of barcoded primers per bead was increased to 10^9^ for sequencing of clone transcriptomes.

To ensure the purity of single-clone encapsulation by CloneSeq, we mixed clones from human and mouse cell lines (PC9 cells and R1 ESCs grown for 7 and 4 days, respectively, in hydrogel spheres) at a concentration of 20,000 and 40,000 clones per ml with flow rates designed to capture one clone per 2 seconds (**Supplementary Table 1**). Both microscopy and a Barnyard (human/mouse) mixing plot of sequencing data showed that at a concentration of 20,000 clones/ml, the encapsulation resulted in excellent separation, with only ~3% of barcodes containing mixed reads, whereas at a higher concentration (>40,000 clones/ml), 10% or more of the reads were mixed (**Figure 2C** and **Supplementary Figure 4**). These result align with a previous analysis of single-cell InDrops, which showed that reducing cell concentration decreased the likelihood of collecting two cells in one drop and decreased the level of contamination of drops with free mRNA^29^.

We next performed scRNA-seq to an average of ~100,000 reads per cell and compared the data to the equivalent sequencing coverage applied for the clones. After reducing PCR duplicates using unique molecular identifier (UMI) counts, we observed that the number of UMIs retrieved from each clone was on average 3 times higher than the numbers retrieved from single cells (average number of UMIs of 30000 for clones and 10000 for single-cells) (**Figure 2D**). The values for the clones were lower than expected by simple clone cell number extrapolations. This is explained by both insufficient primer numbers and reaction inhibition due to the high concentration of cell debris in each drop that originates from the large number of cells composing each clone.

### Impact of 3D hydrogel culture on PC9 cells

To evaluate the impact of our 3D culture method on the cells, we first assessed clone homogeneity. We wished to determine whether the cells within the clone are similar enough to each other to allow us to consider the clone an entity representing the original mother cell. We then compared the expression profile of cells cultured inside the 3D hydrogel spheres to cells grown in 2D to determine if the 3D culture itself altered the cell state. These analyses were performed by leveraging our in-house capability to profile a large number of single cells using drop-based microfluidics^6,31,32^. We produced PC9 cells carrying genomic barcodes located 100 bp upstream to the *BFP* polyA signal. We produced a sequencing library from approximately 50,000 colonies and determined that our plasmid pool contained approximately 20,000 unique barcodes. Barcodes exceeding normal distribution were registered as outcasts for later computational analysis. In these experiments, lentiviruses were transduced into cells with a multiplicity of infection (MOI) of 1 so that only one barcode would enter a cell. We then FACS-sorted BFP-positive cells to collect only virus-containing cells. Next, we encapsulated PC9 cells into 3D hydrogel spheres and grew them until each clone contained 10-30 cells. We then randomly selected about 2000 clones, dissolved the hydrogel spheres to obtain single cells, and performed scRNA-seq on approximately 300 PC9 cells (**Figure 3A**). The limitation of 2000 clones were due to the statistical restraints of the 20,000 clonal barcodes. Cells of the same clone were identified by their identical genomic barcodes. After filtering out cells with unreliable barcodes, we extracted 37 cells originating from 16 distinct clones. The similarity between cell expression profiles was analyzed using tSNE and quantified by calculating the Euclidean distance between points on the tSNE plots as well as the Euclidean distance based on expression profiles^18^ (**Figure 3B**). Euclidean distance between random cells was determined by evaluating all possible cell pairs, and clone distance was calculated as the distance between cells from the same clone. The distance between cells from the same clone was significantly smaller than the distance between randomly selected cells both in the reduction projection space (*p* = 5.96 × 10^−4^) and over the complete transcriptome (*p* = 1.91 × 10^−3^), suggesting that cells of the same clone are indeed significantly more similar to each other than to cells selected at random.

**Figure 3.**
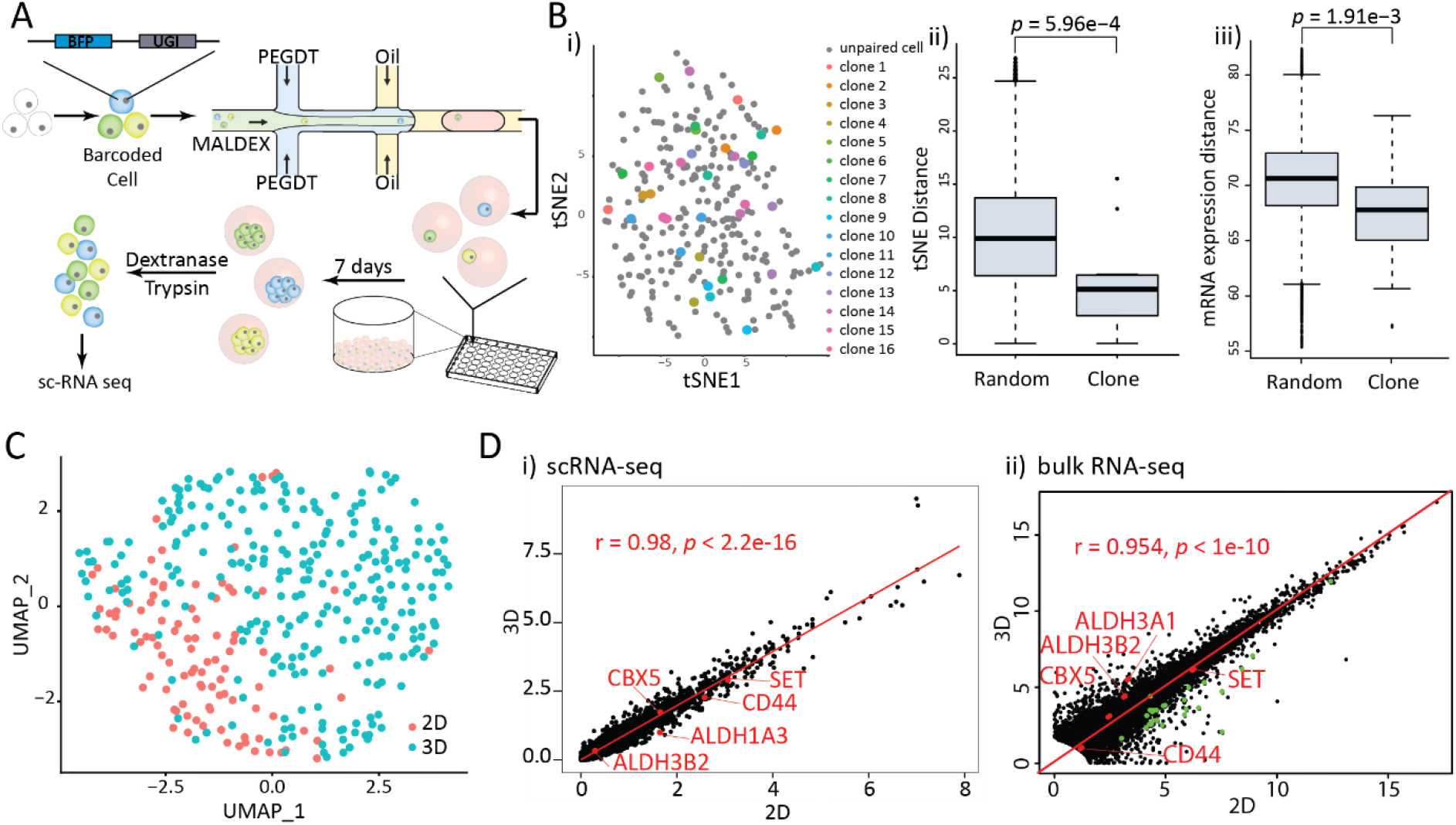
CloneSeq retains single-cell clonal expansion while maintaining cell viability and supports cell “stemness”. (**A**) PC9 cells were gnomically barcoded using lentiviral transfection of 10-basepairs unique guide indices (UGI). Barcoded cells were encapsulated inside the hydrogel spheres and the expanded clones were dissociated, single cells were picked at random, and submitted for scRNA-seq. (**B**) (i) tSNE projection of single-cell transcriptomes of 268 PC9 cells identified 37 paired barcoded cells. (ii) Cell-to-cell tSNE Euclidean distances and (iii) transcriptome Euclidean distances were significantly shorter between cells sharing clonal origin compared with random pairs. (**C**) Single-cell transcriptomes of 272 cells that had been expanded inside 3D hydrogel spheres (3D) and 97 cells that had been expanded in a standard 2D culture (2D) were mapped into one cluster on a UMAP space. (**D**) A correlation between the gene expression levels of cells cultured in standard 2D and in 3D configurations is maintained both via (i) scRNA-seq and (ii) bulk RNA-seq. Gene levels were averaged across single cell transcriptomes. High correlation excludes a significant 3D culture effect. Bulk RNA-seq sensitivity reveals stemness signature that are supported by 3D cultures (red dots) and cell cycle signatures in 2D (green dots). R: Pearson coefficient of correlation.

PC9 cells grown in standard 2D conditions have a doubling time of ~1.5 days^33^. In our 3D spheres, PC9 cells were less proliferative, with a doubling time of ~2.5 days. To further evaluate the impact of 3D culture on the cells, we compared PC9 cells growing in 2D versus 3D conditions, using both scRNA-seq and bulk RNA-seq. Single cell-based UMAP projections did not capture substantial differences between the 2D and the 3D culture conditions (**Figure 3C**). In support, gene expression comparison of scRNA-seq showed a highly significant correlation coefficient (0.98; *P* < 10^−16^), similar to the overall correlation observed for bulk RNA-seq (0.95; *P* < 10^−10^) but the latter also revealed the upregulation of cell cycle-related genes in 2D (green dots, **Figure 3D**). In contrast, bulk RNA-seq revealed that 3D culture led to the induction of genes associated with adhesiveness and cancer stem cell like signature^34^ (e.g. *Aldh3A1* and *Aldh3A2*, **Figure 3D**). Other genes known to be associated with CSC signature (*SET, CBX5 and CD44*, **Figure 3D**) were found equally expressed in both 2D and 3D culture conditions. As these genes are not highly expressed in PC9 cells, they couldn’t be detected by scRNA-seq. Taken together, these data suggest that while cells grown in 3D culture in PEGDT-MALDEX hydrogel are overall similar to those grown in 2D conditions, there are differences in expression of some genes associated with, e.g., cancer stemness in PC9 cells.

### Clone-to-clone variation identifies novel subpopulations

To study the effect that clonal expansion within the hydrogel spheres has on the homogeneity of cell states within the clones, we compared the inter-clone correlations of small (n<15 cells) and large (n≥15 cells) clones and of pseudo-clones that were formed *in silico* by averaging randomly sampled single-cell transcriptomes. As expected, the correlation between single-cell transcriptomes that were randomly sampled from a heterogeneous population of PC9 cell states is low, thus reflecting the inherent degree of cell-to-cell variation and limited complexity (Fig. 4A). With increasing cell number, cell-to-cell variations are averaged. Hence, the transcriptional correlation between “pseudo-clones” increased with cell number plateauing at n=8 cells (Fig. 4A). Consistently, the correlation between the transcriptomes of small PC9 clones (<15 cells) was lower than between large PC9 clones (n≥15 cells). However, clone-to-clone transcriptional correlation of both small and large clones was comparable with 2-cells pseudo-clones and significantly lower than pseudo-clones of similar size. This indicates that clonal expansion within the hydrogel spheres maintains cell state and hinders cell-to-cell variation compared with the transcriptomes of non-clonal cells. Importantly, this variation is not the outcome of cell cycle states or transcriptional bursting, as those are averaged in clones. In this manner, CloneSeq amplifies sequencing sensitivity and coverage by reading the transcripts that were pooled from multiple cells while excluding averaging out distinct cellular states.

**Figure 4.**
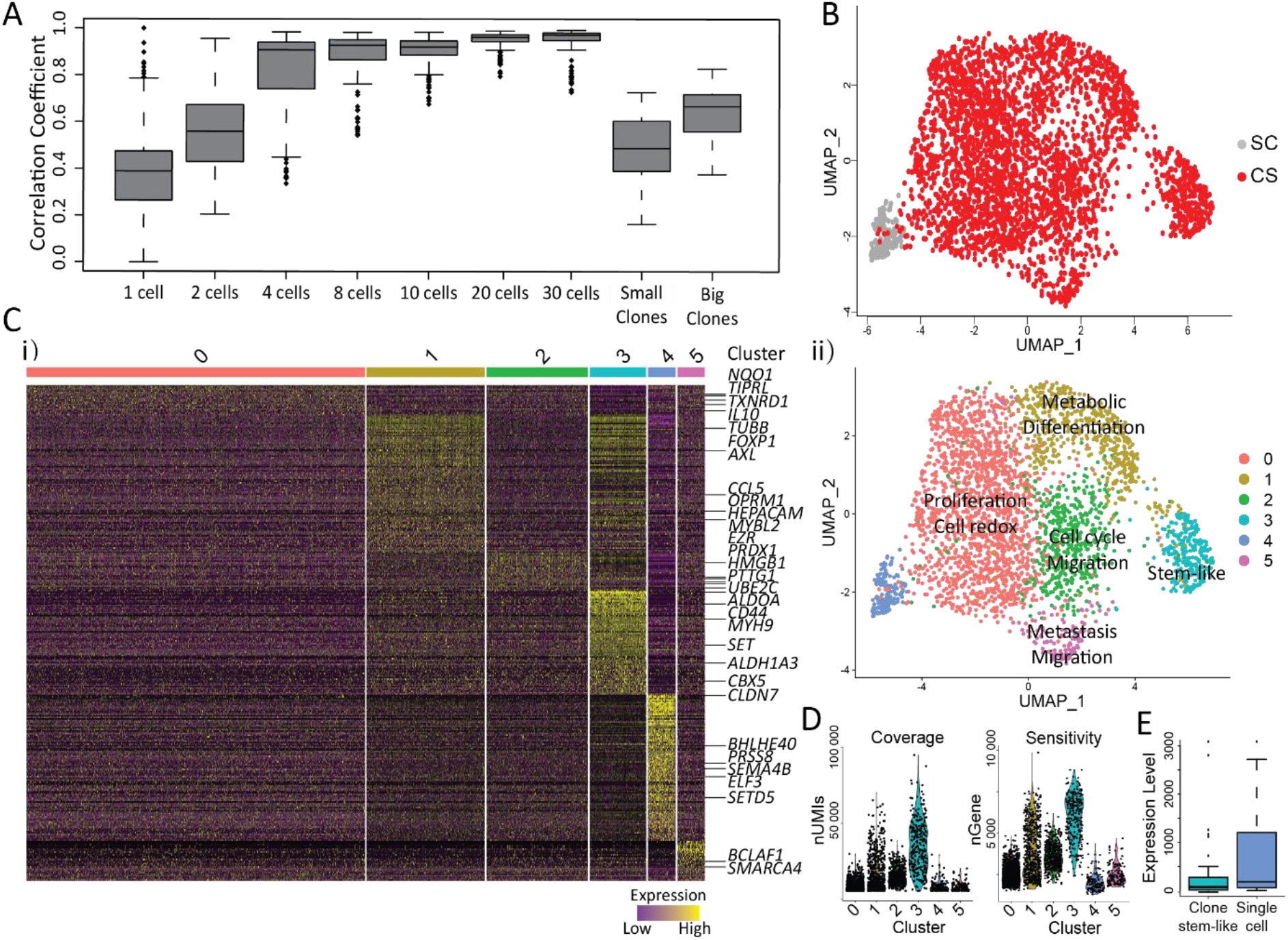
Clone-to-clone variation analysis for PC9 cells. (**A**) The transcriptional correlation between pseudo-clones that were *in silico* aggregated increases due to averaging out of pseudo-clone to pseudo-clone variation with increasing pseudo-clone size, plateauing at n = 8 cells, while the correlation among real clones remains relatively low and clone-to-clone variation is not averaged out both for small clones (n ≤ 15 cells) and for big clones (n > 15 cells). (**B**) Clones at day-7 after encapsulation were transcriptionally profiled either after dissociation into single cells (SC: scRNA-seq; n = 300, UMI>5000) or as clones (CS: CloneSeq; n = 3000, UMI>10000) and mapped onto UMAP space. (**C**) (i) Most differentially expressed genes are identified via KNN unsupervised clustering of scRNA and CloneSeq transcriptomes, (ii) underlying distinctive MSigDB GO term and cell function annotations. (**D**) The expression levels of the marker genes of single cell cluster-4 are highly expressed compared with the marker genes of the stem-like cluster-3 (p < 0.01). (**E**) Compared with CloneSeq, scRNA transcriptome show low transcript sensitivity (nUMI) and low gene coverage (nGene).

To evaluate the association between single-clone transcriptomes and single-cell transcriptomes, we cultured PC9 clones for 7 days and performed Clone-Seq for 4000 clones and 300 single cells dissociated from clones. Single-clone and single-cell transcriptomes were dimensionally-reduced and clustered via *k*-nearest neighbors and shared nearest neighbor (SNN) graph clustering. Six clusters were identified and projected onto a dimensionally-reduced UMAP space (Seurat package 3.0; Fig. 4B-i,ii).^35,36^ Single-cell transcriptomes were clustered into cluster-4 and separated from single-clone transcriptomes that were divided between five clusters. To test the option that transcriptome clustering was dominantly driven by clone size, we down-sampled both CloneSeq and single-cell transcriptomes to 5000 UMIs. Despite this effective reduction in transcript detection, we successfully recapitulated the separation between transcriptome clustering (**Supplementary Figure 5A)** as well as between single-cell and single-clone transcriptomes (**Supplementary Figure 5B)**. UMI downsampling analysis thus indicates that CloneSeq provides superior sensitivity enable cell-state based clustering independent of direct clone coverage.

Cluster identities were characterized using DisGeNET^37^ and MSigDB^38^. Overall, 300 differentially expressed genes explained the different clusters for which 46 were previously known as NSCLC related (Supplementary Table 3). Cluster 4 was marked by highly expressed genes with mixed signatures (Fig. 4C). For example, *ULK1* promotes cell proliferation, whereas *BHLHE40* inhibits proliferation and induces cellular senescence; *SETD5* stimulates whereas *CLDN7* inhibits tumor invasion and migration; *ELF3* supports whereas *SEMA4B* and *PRSS8* suppress tumor growth and metastasis. Unlike cluster-4, the remaining five clusters that were characterized by known genes of non-small cell lung cancer (NSCLC) were distinct in their functions. Cluster-specific marker genes and functions are summarized in **Table 1**. Cluster-0 was marked by genes associated with cancer cell progression and proliferation, redox genes, and drug response genes. Cluster-1 was enriched for genes involved in RNA metabolism and cell differentiation – specifically towards the myeloid axis. Interestingly, Cluster 1 was marked by *AXL* and *OPRM1* that promote tumor growth, and by *FOXP1* that is implicated in epithelial to mesenchymal transition (EMT).^39–41^ Cluster 2 was enriched by genes that regulate cell migration, proliferation, and cell cycle, including *UBE2C*, *PTTG1*, *HMGB1*, *PRDX1*, *EZR*, *and MYBL2.* Genes marking Cluster 3 (e.g. *CBX5, ALDH1A3*, *SET* and *CD44*) are strongly related to CSCs^34,42^. Cluster 5 is the smallest one, consisting of clones that are marked by genes associated with metastasis and migration (*RAC1*, *PPIA*, *BCLAF1*).

**Table 1.**
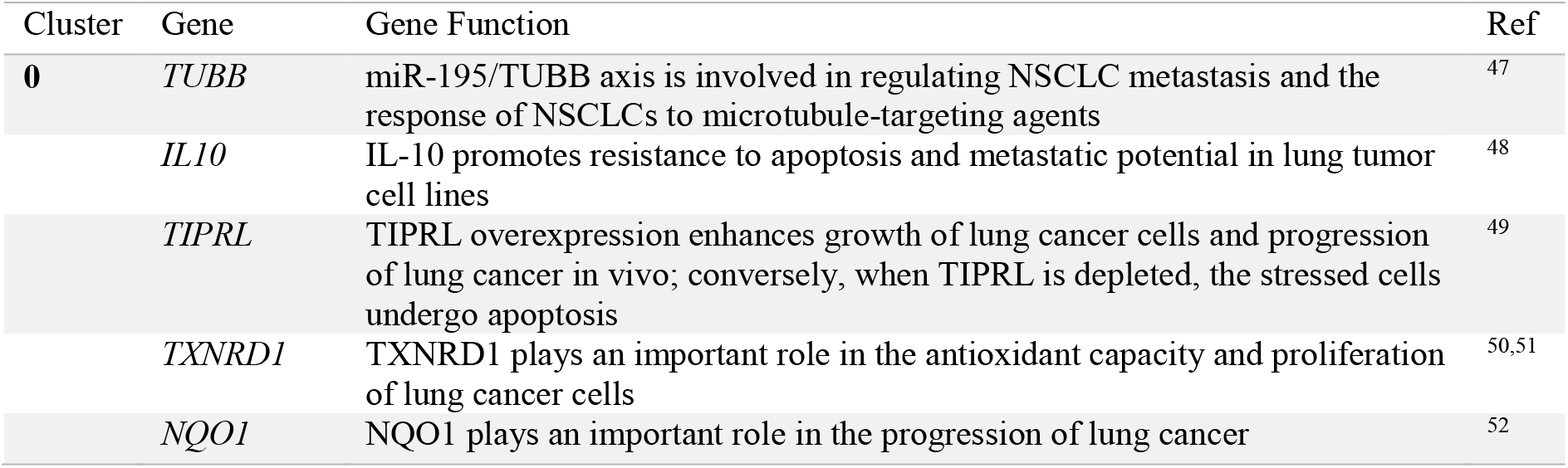
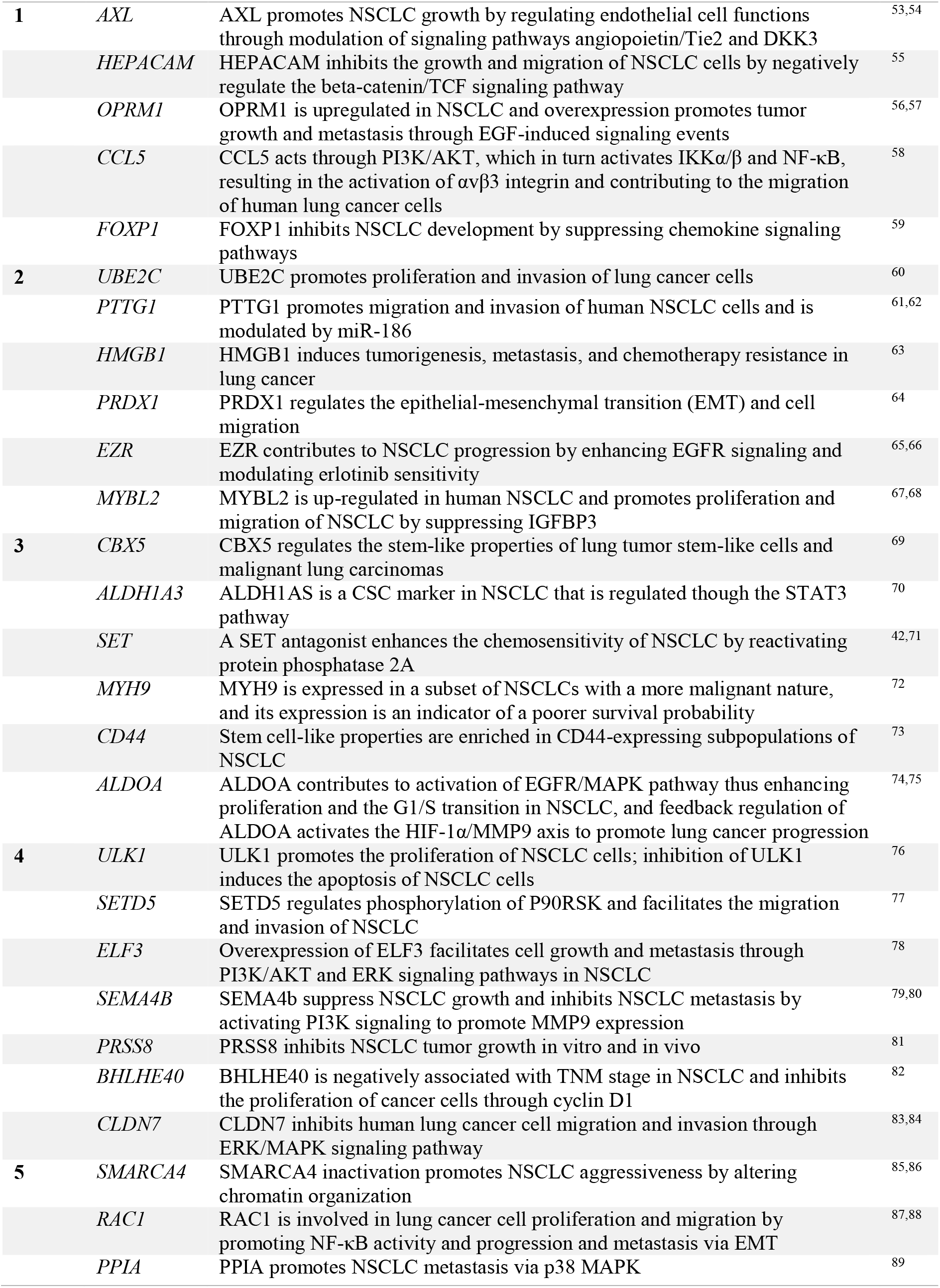
Representative genes enrichment in clusters of PC9 single cells and clones

The transcriptional coverage and sensitivity detected by CloneSeq were both higher than by single-cell RNA-seq (cluster-4; Fig. 1D). In particular, cluster-3 consists of the largest clones that maximized both transcript number and gene coverage, which is in line with the in vitro proliferation potential of CSCs^43–45^. Herreros-Pomares *et al.* previously showed that NSCLC CSCs are highly proliferative in sphere-forming culture^46^. Interestingly, cluster-specific differences in transcriptional sensitivity and coverage were reproduced even for down-sampled transcriptomes (Fig. S6). We compared gene expression levels of marker genes of cluster-4 (single-cell transcriptomes) and cluster-3 (CSC-like clones) using unbiased bulk RNA-seq. We found that Cluster-4 marker genes were expressed at higher levels compared with the marker genes of Cluster-3 (**Figure 4E**). Together, we conclude that the improved sensitivity that is provided by CloneSeq enables the detection of low-to-mid expressed genes, which fail to be detected via single-cell RNA-seq. Since cell-states are often characterized not only by highly-expressed marker genes but also by lowly expressed genes (e.g. regulatory genes), CloseSeq provides means for elucidating the existence of distinct cell states within a heterogenous population of cells, which is not accommodated by single-cell RNA-seq methods.

### 3D hydrogel spheres promote stemness

As our 3D culturing method induced proliferation of CSCs, we hypothesized that the 3D conditions within the gels promote stemness. To test this, we explored the effect of 3D culturing on ESCs. Ground-state pluripotency can be maintained *in vitro* by culturing the cells with LIF together with GSK3/MEK inhibitors (‘2i’)^90,91^. The microenvironment in which the cells grow influences the cell state and can activate or repress differentiation pathways^92^. ESCs grown without 2i-LIF do not retain pluripotency and they differentiate spontaneously along the different lineages (ectoderm, endoderm, mesoderm, and extraembryonic endoderm)^90^.

To validate the effect of the 3D environment on stemness, we used ESCs that express three pluripotency reporters: GFP-NANOG and BFP-ESRRB as markers for ground-state pluripotency and RFP-UTF1 for the primed state^93–95^. As expected, ESCs grown in both our 3D system and the 2D gold standard (0.1% gelatin) setup in the presence of 2i-LIF expressed all three pluripotent markers (**Figure 5A**). However, when the cells were grown without 2i and LIF for 6 days, cells in the 2D system quickly lost their pluripotency markers, whereas cells growing in 3D continued to express these markers, with NANOG and ESRRB slightly down-regulated, and UTF1 slightly upregulated (**Figure 5A**). This suggests that our 3D hydrogel spheres support pluripotency and significantly delay ESC spontaneous differentiation in the absence of 2i-LIF, possibly explaining the highly homogenous states observed for the ESC clones grown in 3D. To further validate the pluripotency of the cells after 8 days in the absence of 2i-LIF, the hydrogel spheres were removed, and the cells were re-seeded on mouse embryonic fibroblasts (MEFs) for 4 days. In cells originating from the hydrogel spheres, about 70% (average of two independent experiments) of the cells formed Nanog-positive colonies, whereas less than 8% of the cells originating from the 2D condition formed Nanog-positive colonies (**Figure 5B**). To corroborate these findings, we performed RNA-seq on cells grown in 2D and 3D conditions for 4 days without 2i-LIF. The results confirm higher expression level of pluripotent transcription factors in cells grown in hydrogel spheres compared to cells grown on gelatin (**Figure 5C**). This demonstrates that the 3D hydrogel spheres help maintain a pluripotent state.

**Figure 5.**
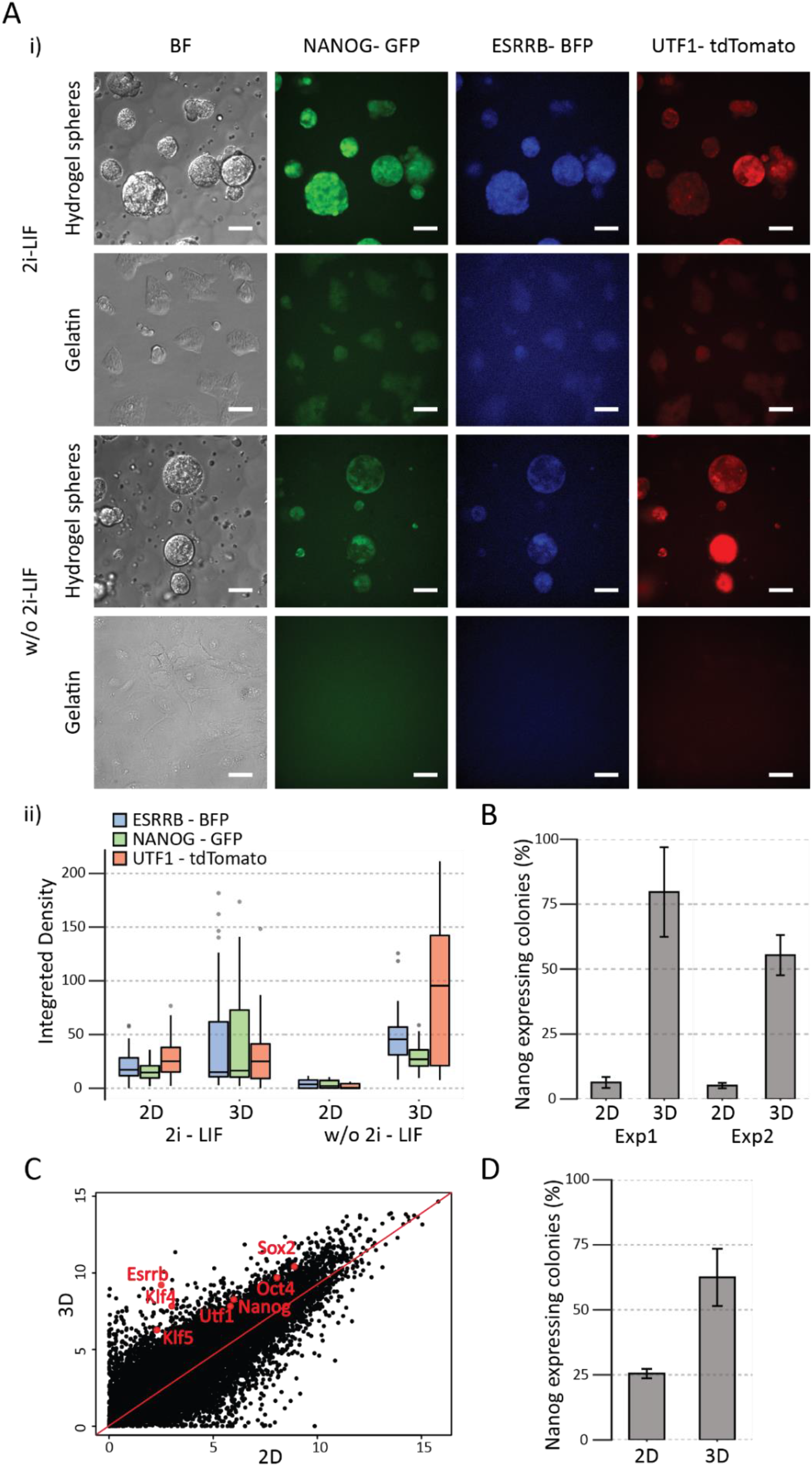
3D soft hydrogels support a pluripotent state even without 2i-LIF. (**A**) i) Representative NANOG, ESRRB, and UTF1 immunofluorescence staining of 3D ESC clones encapsulated inside hydrogel spheres and 2D ESC colonies cultured on gelatin-coated plates. Cells were cultured for six days with or without 2i-LIF supporting factors. Scale bars, 50μm. ii) Immunofluorescence density of NANOG and ESRRB is maintained within the hydrogels spheres even without 2i-LIF supporting factors whereas UTF1 levels increase (average of 30 colonies per condition). (**B**) The percent of NANOG-positive colonies are compared between ESCs that were cultured inside 3D soft hydrogel spheres and on gelatin-coated 2D plates for 8 days, dissociated and further cultured on MEF feeder layer for subsequent 4 days (average of three representative regions in the plate). *P-value* < 0.001. (**C**) The expression of pluripotent genes (marked in red) is higher in cells that were cultured inside 3D hydrogel spheres (3D) compared with cells that were cultured on gelatin-coated plates (2D) without supporting factors. (**D**) MEFs were encapsulated inside hydrogel soft spheres (3D) or seeded on gelatin-coated plates (2D) and epigenetically reprogrammed using tetracycline-activated OSKM cassette. After 10 days of reprogramming, cells were dissociated, seeded onto MEF feeder layer and the fraction of NANOG-positive colonies was compared (average of three areas in the plate).

Finally, we tested whether the 3D microenvironment enhances the reprogramming efficiency of MEFs into induced pluripotent stem cells (iPSCs). We compared MEFs containing the OSKM cassette under the control of the TET-on promoter^96^ grown in standard iPSC conditions in 2D, to the same MEFs grown inside our 3D hydrogel system (**Supplementary Figure 6**). After 10 days, during which the cells were supplemented with 4.5 μM tetracycline, we seeded the cells on MEFs and quantified Nanog positive colonies. Reassuringly, the 3D culture showed ~2.4-fold higher frequency of Nanog-positive cells compared to the 2D cultures (**Figure 5D**). One clear disadvantage of reprogramming in the 3D setup is that reprogramming is accompanied by a high number of dead cells that failed to complete their cellular transformation^97^. These apoptotic cells accumulate inside the hydrogel spheres and cannot be washed away easily. As apoptotic cells secrete signals that interfere with iPSC formation, we predict that iPSC production efficiency could be significantly improved by adding cycles of breaking down hydrogel spheres, removing dead cells, and re-encapsulating MEFs.

## Discussion

Single-cell based assays have revealed the scope and importance of heterogeneity in many biological systems^98–103^. However, single-cell sequencing technologies are of low sensitivity, and therefore they are prone to detect cell-to-cell variation based on highly expressed genes while missing differences in expression of low-abundance genes^104^. In addition, the influence of stochastic effects, such as transcriptional bursting and cell-cycle state, on single-cell data can potentially mask important aspects of cellular heterogeneity. To overcome this obstacle, we developed an innovative technology that allows single cells to be grown into small clones in 3D hydrogel spheres.

The 3D culturing system is based on microfluidics and uses a PEGDT-MALDEX-based biodegradable hydrogel spheres. The hydrogel spheres mimics the characteristics of natural extracellular matrices. Since we used a biodegradable component (dextran), the hydrogel spheres can be dissolved in bulk or as microfluidic droplets. Our results suggest that the biochemical composition and the physical structure of the hydrogel spheres are stable, which ensures the compartmentalization of single cells during their development into clones. To date, we have optimized 3D hydrogel sphere culture conditions for growth of PC9 human lung cancer cells as well as mouse ESCs and MEFs that were reprogrammed.

The ability to profile small clones improved sensitivity of detection of subtle cellular states when compared to single-cell analysis, as it minimizes the presence of confounders such as cell cycle and transcriptional bursting, and because it supports the capture of a greater number of transcripts per clone compared to single cells. We showed for human PC9 cells that cells that share a clonal origin are more similar to each other compared to random cells, suggesting that clones are homogeneous. This will make clonal-based profiling a valuable tool that can potentially improve the low sensitivity of standard single-cell assays.

Cancer cells didn’t grow with the same speed inside hydrogel spheres, with PC9 clone sizes ranging from 1 to 30 cells after 7 days of culture. We verified that the undivided single cells were alive by trypan blue staining. Slowly replicating cells had different gene expression signatures from rapidly dividing cells in our single-clone analysis. Differentially expressed genes between the two were associated with cellular migration, growth inhibition, and metastatic state. Interestingly, some of the clones with higher numbers of cells were also associated with a cancer stem cell-like signature. As our single-cell reference profiles were based on cells growing in 3D, and these cells did not show significantly different cellular states compared to single cells grown in regular 2D conditions, we suggest that the stem cell-like state, at least in PC9 cells, cannot be detected with standard single-cell profiling. Moreover, while CSCs are rare in the body^105^ they can be enriched by sphere-forming culture^46^. Our analyses of PC9 cells indicated that the 3D culture system promotes stemness, and we further validated this by culturing and analyzing ESCs using CloneSeq.

When grown in PEGDT-MALDEX hydrogel spheres for 8 days, ESCs maintained pluripotency even in the absence of 2i/LIF. Moreover, 70% of the ESCs were pluripotent after release from the hydrogel spheres and seeding on MEFs for 4 days, whereas only 8% of the cells originating from the 2D condition were pluripotent. RNA-seq results corroborated our findings that cells grown in 3D conditions showed significantly higher expression of pluripotent transcription factors. Together, these results indicate that our 3D culturing system promotes stemness.

In summary, CloneSeq is an effective and general method that leverages 3D culture, drop-based microfluidics, barcoding, and high-throughput sequencing to dramatically extend our ability to characterize cellular states with great statistical power. We plan to optimize CloneSeq to support culturing of primary tumor-derived cells to allow high sensitivity, clonal profiling of cancer cellular states in association with different treatments. In addition, the method could be used to enable full-length RNA-seq to dissect alternative splicing events and somatic mutations in cancer samples. ChIP-seq and bisulfite-seq protocols could also be performed on clones grown in the hydrogel spheres. We expect that CloneSeq will expand our understanding of cancer biology in particular and of other biological systems involving proliferative cells.

## Acknowledgments

O.R. is supported by research grants from the European Research Council (ERC, # 715260 SC-EpiCode), the Israeli Center of Research Excellence (I-CORE) program, the Israel Science Foundation (ISF, #1618/16), and Azriely Foundation Scholar Program for Distinguished Junior Faculty. O.R and A.A are supported by Nofar (65883) of the Israel Innovation authority. E.M. is the Arthur Gutterman Family Chair in Stem Cell Research and is supported by the Israel Science Foundation (ISF1140/17). This project has received funding from the European Union’s Horizon 2020 research and innovation programme under the Marie Skłodowska-Curie grant agreement No 765966 - EpiSyStem.

## Author Contributions

D.B., X.S., C.K., D.E, E.M, A.B. and O.R. conceived the study, prepared the figures and wrote the manuscript. D.B., X.S., E.M, A.B, and O.R. designed the experiments. C.K, A.A. A.M, A.Bi and D.E. performed PC9, ESCs and IPSCs tissue culture, and library preparation. S.B and D.B preformed microscopy and Pipet aspiration test. D.B. and X.S. prepared the microfluidics system and performed scRNA-seq and CloneSeq experiments. X.S and C.K preformed clonal barcoding experiments. X.S., D.E and O.R. preformed the computational analysis.

## Author Information

All bulk, single-cell RNA-seq and CloneSeq data will be deposited in the Gene Expression Omnibus database (GEO). Reviewers, please use this link: in process. The authors declare no competing financial interests. Correspondence and requests for materials should be addressed to A.B and O.R.

## Materials and methods

### Cell culture

PC9 lung adenocarcinoma cells that express GFP and R1 mouse embryonic stem cells were kindly provided by Prof. Ravid Straussman (Weizmann Institute, Israel). BYKE1 and OSKM 2nd MEFs cells were kindly provided by Dr. Yosef Buganim (The Hebrew University, Jerusalem). PC9 cells were grown in DMEM (Sigma-Aldrich, cat #D5671) supplemented with 10% fetal bovine serum (FBS; Biological Industries Israel, cat #04-007-1A), 50 μg/ml penicillin-streptomycin (Biological Industries Israel, cat #03-031-1B), 2 mM L-glutamine (Biological Industries Israel, cat #03-020-1B), and 1 mM sodium pyruvate (Biological Industries Israel, cat #03-042-1B). R1 and BYKE1 ESCs cells were grown on 0.1% gelatin-coated standard tissue culture dishes and maintained in ESC medium (DMEM, 15% ESC-grade FBS, 50 μg/ml penicillin-streptomycin, 2 mM L-glutamine, 1 mM sodium pyruvate, 0.1 mM non-essential amino acids (Biological Industries Israel, cat #01-340-1B), 0.1 mM β-mercaptoethanol (Sigma-Aldrich, cat #M3148)). To maintain pluripotency, 1000 U/ml LIF and 2i (1 μM PD0325901, 3 μM CHIR99021) were added to the culture medium. OSKM MEFs were grown on 0.1% gelatin in DMEM together with 10% FBS, 50 μg/ml penicillin, 2 mM L-glutamine, 1 mM sodium pyruvate, and 0.1 mM non-essential amino acids. For differentiation experiments, cells or hydrogel spheres were washed with PBS, and resuspended in basal ESC medium 1) without 2i, 2) without 2i and LIF, or and 3) without 2i and LIF supplemented with 0.25 M all-*trans*-retinoic acid (Sigma-Aldrich, Cat #R2625), and cultured for 4 days.

### Device design and fabrication

Four microfluidics devices were used: (I) to produce acrylamide hydrogel microparticles with acrydite-modified DNA primers for barcoding; (II) to encapsulate single cells within PEGDT/MALDEX hydrogels; (III) to encapsulate single cells with barcodes, lysis buffer, and reverse transcriptase (RT) enzyme within droplets; and (IV) to encapsulate clones with barcodes, lysis buffer, and RT enzyme within droplets (**Supplementary Figure 3A-D**). Devices were designed using the AutoCAD software (Autodesk). All chips were fabricated by photolithographically defining SU8 (SU-8 2050, MicroChem) on silicon wafers at the Harvey Krueger Center of Nanotechnology at the Hebrew University of Jerusalem. The depths of the photoresist layers were 50.4 ± 1 μm for device I, 81.2 ± 1 μm for device II, 79.1 ± 1 μm for device III, and 123.6 μm ± 1 μm for device IV. Designs for devices I and III were adapted from inDrops^29^. Device II was modified from the co-flow drop maker of Hi-SCL^32^, whereas device IV is a modification of the standard inDrops chip. The designs used to fabricate the devices are available in CAD format (**Supplementary Files**). Polydimethylsiloxane (PDMS) at a ratio of 10:1 of base and crosslinker, respectively, was formed by curing the prepolymer (Sylgard 184, Dow-Corning) on the silicon templet at 65 °C for 2 h. PDMS devices were covalently bound to N°1 glass coverslips using Femto oxygen plasma activation (Diener) for 15 s at 50 W. The PDMS devices were treated with Aquapel (Rider) water repellent and dried under air in order to make the devices more hydrophobic and prevent wetting of drops on the channel walls. Drop volume (*v*) calculation for CloneSeq devices was based on still images of droplets at the outlet of the microfluidic device using the equation: 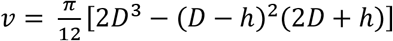, where *h* is the height of the channel and *D* is the droplet diameter in μm^29^.

### Barcode and primer design

The acrydite-modified DNA primers used for the hydrogel barcode beads are based on a previously published protocol29; primers were supplied by IDT. We modified the barcode plates (eight 96-well plates) to expand barcode complexity by changing the length of barcode 1 to a variable length of 7 to 10 bases. DNA oligonucleotide sequences are listed in the Supplementary File. We typically used a 10-nmol scale normalization and standard desalting, and ordered oligonucleotides dissolved to a final concentration of 50 μM in 10 mM Tris-HCl (pH 8.0), 0.1 mM EDTA (TE buffer).

### Production of hydrogel beads carrying barcoded oligonucleotide primers

Hydrogel beads carrying barcoded DNA primers were produced using a method described previously^29^. The hydrogel beads were composed of a 4xAB solution (2.6 ml 40% acrylamide in water (Sigma-Aldrich, cat #01697), 3.6 ml 40% total 19:1 acrylamide:bis-acrylamide aqueous solution (Bio-lab, cat #1352335), 3.8 ml water) supplemented with an acrydite-modified DNA primer (5’-ACryd/iSpPC/CGATGACGTAATACGACTCACTATAGGGATACCACCATGGCTCTTTCCCTACACGACGCTC TTC-3’, where Acryd represents acrydite and iSpPC represents the photo-cleavable Int PC). DNA primers on polymerized hydrogel beads were barcoded using a combination of the split-and-pool method and a primer extension reaction using an automated liquid handling system (Biomek 4000, Beckman Coulter Life Science). The final barcode library complexity was around 147,456 unique barcodes repeated across around 40 million hydrogel beads per synthesis batch with an average of 10^9^ copies of fully extended DNA primers per single bead^29^. Hydrogel beads were produced using the flow-focusing microfluidic device I (**Supplementary Figure 3A**) as previously described^106^. Flow rates used for the hydrogel bead synthesis were 1000 μl/h for a 4xAB solution supplemented with an acrydite-modified DNA primer and 1600 μl/h for the oil phase (**Supplementary Table 1**). After barcode synthesis, the barcoded beads were filtered twice with a cell strainer of 70 μm (pluriSelect, Cat #43-10070-50) to obtain homogeneously sized beads with a diameter of around 60 μm, as shown in **Supplementary Figure 3E**.

### Microfluidics operation

Droplet formation and cell encapsulation were performed using EZ pressure-derived pumps (Fluigent), controlled by the A-I-O software at pressures ranged from 69 mbar to 2 bar. The continuous oil phase for all droplet microfluidics experiments was Novec HFE-7500 fluorinated oil (3M) containing 2% w/w 008-FluoroSurfactant (RAN Biotechnologies). For all experiments, cells were kept in a tube surrounded by ice and were gently agitated with a micro-stir bar placed inside the tube and rotated using a magnet attached to a rotating motor to prevent sedimentation and clumping. The flow was visualized under an optical microscope (NIKON Ti-U) at 10x magnification and imaged at 1000-2000 frames per second using a Hispec1 camera (FASTEC Imaging). **Supplementary Table 1** summarizes the flow rates and the pressure pumps used for operating the different devices used.

**Supplementary Table 1.**
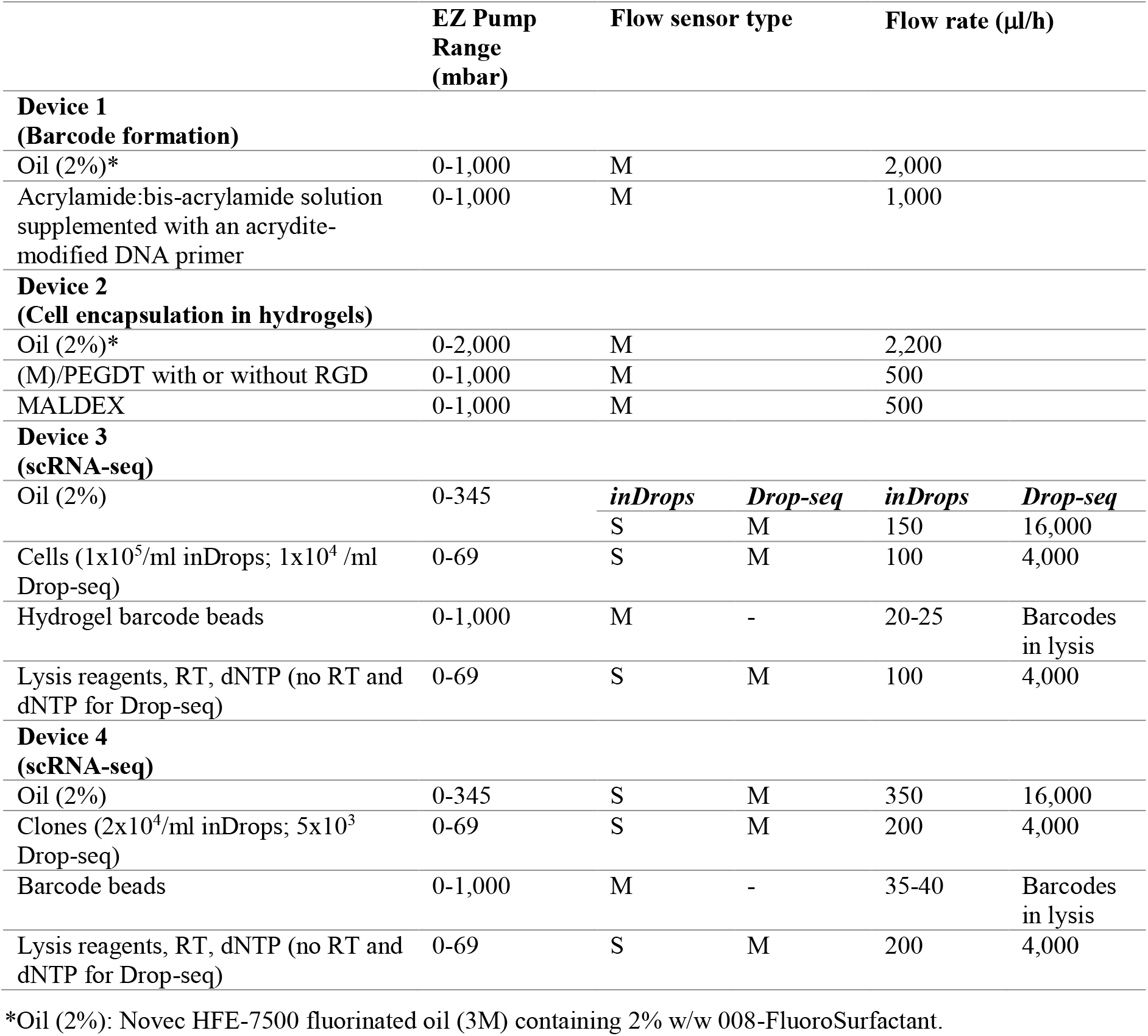
Flow rate and pressure pumps used for operating the different devices.

### Clone formation within PEGDT/MALDEX hydrogels

All materials used to synthesize and dissolve different hydrogel spheres used to grow PC9 and ESCs cells were from Cellendes GmbH, Germany. We used PEGDT (Mn ~10,000; cat #L50-1) and MALDEX (cat #M92-3) hydrogel chemistry to encapsulate cells within spheres and grew them into clones. PC9 cells were encapsulated in PEGDT/MALDEX hydrogel with no additional cell adhesion peptides or remodeling supplements. ESCs cells were encapsulated in MALDEX and cell adhesion peptides containing cell recognition motifs of the extracellular matrix (RGD (cat #P10-3); peptide sequence: Acetyl-Cys-Doa*-Doa-Gly-Arg-Gly-Asp-Ser-Pro-NH2 [*Doa:8-amino-3,6-dioxaoctanoic acid]) and MMP-cleavable peptide modified PEGDT (cat #L60-1; MMP sequence: Pro-Leu-Gly-Leu-Trp-Ala). Hydrogel spheres were dissolved using a 1:20 dilution of dextranase from *Chaetomium gracile* (cat #D10-1) in PBS incubated for 30 min at 37 °C. The gelation buffer (GB; cat #B20-3) used in all cell encapsulations contained 10 g/l glucose, 0.5 M HEPES (pH 7.2), 0.05 M KCl, 1.1 M NaCl, 0.2 M NaH_2_PO_4_, and 0.2 g/l phenol red. Before use, MALDEX, PEGDT, MPEGDT, and RGD peptide were briefly spun down to make sure that the lipolysis material was at the bottom of the reaction tube. MALDEX was resuspended in 170 μl of doubly distilled water to a concentration of 30 mM maleimide groups. PEGDT and MPEGDT were resuspended in 188 μl of doubly distilled water to a concentration of 20 mM thiol groups. RGD peptide was resuspended in 48 μl of doubly distilled water to a concentration of 20 mM of peptide and thiol groups.

Device II (**Supplementary Figure 3B**) was used to encapsulate the single cells within the hydrogel spheres. For PC9 cells, inlet 1 consisted of 310 μl 0.1% w/v gelatin in doubly distilled water, 50 μl of GB, 67.5 μl of 20 mM PEGDT, and 120 μl PBS. For R1 ES cells, inlet 1 consisted of 300 μl 0.1% w/v gelatin in water, 10 μl of 20 mM RGD peptide, 50 μl of GB, 67.5 μl of 20 mM MPEGDT, and 120 μl PBS. For both cell types, inlet 2 consisted of 335 μl 0.1% w/v gelatin in water, 50 μl of GB, 45 μl of 30 mM MALDEX, and 120 μl of cell suspension containing around 1 million cells in 99 μl PBS, 10 μl Extracellular matrix (Sigma-Aldrich, cat #E1270), and 11 μl OptiPrep Density Gradient Medium (Sigma-Aldrich, cat #D1556) to minimize cell clumping. The resulting hydrogel-cell mix was subsequently enveloped in the device in HFE 7500 oil with 2% surfactant (inlet 3) to produce single-cell hydrogel spheres of 50-55 μm in diameter. Flow rates were 500 μl/h for inlets 1 and 2 and 2200 μl/h for inlet 3 (**Supplementary Table 1**). The resulting single cell-containing hydrogel spheres were allowed to cure for 5 min at 37 °C, and the upper hydrogel fraction (~500 μl) was demulsified by incubating 1 min at 37 °C in the demulsifying solution containing 400 μl of neat HFE 7500 oil, 100 μl perfluoro-1-octanol (PFO, Sigma-Aldrich, cat #370533), 280 μl PBS, and 20 μl of 1 g/ml methoxy PEG thiol (average Mn ~800; Sigma-Aldrich, cat #729108). Methoxy PEG thiol was used to mask unbound maleimide groups and to prevent aggregation of spheres while demulsifying. The hydrogel spheres fraction was washed three times with 1 ml PBS with centrifugation at 250 rcf for 2 min. For culturing encapsulated cells into clones, 500-μl aliquots of hydrogel spheres were grown in a standard 48-well cell culture dish well with 500 μl standard cell growth medium suitable for each cell type and growth condition.

### Viability evaluation of clones within hydrogels

Cell viability within PEGDT/MALDEX hydrogels was evaluated by counting cells stained with trypan blue (Biological Industries, Israel, cat #03-102-1B). Cells were encapsulated in PEGDT/MALDEX based hydrogels as single cells and then left to grow into clones for 4 days. The same culture media were used as for normal cell tissue culture. The media was replaced daily for ESCs and every 3 days for PC9 cells. After 4 days, hydrogel spheres were washed three times with 1 ml PBS with centrifugation at 250 rcf for 2 min. Beads were then suspended in 300 μl of 1:20 dilution of dextranase from *Chaetomium gracile* in PBS and incubated for 30 min at 37 °C. Following hydrogel spheres degradation, cells were centrifuged at 500 rcf and treated with 100 μl 1x trypsin/EDTA solution for 5 min at 37 °C in order to break aggregates. The trypsin was then quenched by adding an equal volume of medium. Trypan blue was added at a 1:1 ratio with the cell medium. Live/dead cell viability was assessed using Countess II FL Automated Cell Counters (Thermo Fisher Scientific). For each condition, cell counts were obtained for three different measurements averaged over two batches replicates.

### Hydrogel spheres microenvironment and mechanical properties

To evaluate the microenvironment in hydrogel spheres, we used GFP-based R1 ESCs and PC9 cells encapsulated in rhodamine-modified PEGDT/MALDEX hydrogel spheres. To modify the spheres, we added 12.5 μl of 1 mg/ml of Biotin PEG thiol, MW 400 (NANOCS, cat #PG2-BNTH-400) to inlet 1, which contained the PEGDT when encapsulating single cells within the hydrogel spheres. Following sphere formation, the biotin-modified hydrogel spheres were washed three times with PBS, modified with a 1:1 volume ratio of packed hydrogel spheres and 1 mg/ml streptavidin-rhodamine (Jackson ImmunoResearch, Cat # 016-290-084) for 5 min, and washed three times with PBS.

To analyze sphere homogeneity, empty biotin modified hydrogel spheres were modified with 1:1 volume ratio of 1 mg/ml streptavidin (Sigma-Aldrich, cat #85878) for 5 min, washed three times with PBS, and then modified with 1:1 volume ratio with 1 mg/ml biotin-5-fluorescein conjugate (Sigma-Aldrich, cat #53608) followed by three washes with PBS. Cross-sectional images were taken using NIKON A1 confocal microscope to evaluate the distribution of molecules within the hydrogel spheres.

Hydrogel spheres mechanics were measured using a micropipette aspiration system made in-house using a micromanipulator holder (Narishige, NT-88-V3) connected to a manual hydraulic, oil-filled microinjector (Eppendorf, CellTram) and a pressure-sensing diaphragm (Validyne, DP15, CD379). Micropipettes were fabricated by pulling borosilicate capillaries (Sutter Instruments, MicroPuller P-1000) and then forging them to 3-μm inner diameter tips (Narishige, MF-830 Micro-Forger). Suspended beads were then placed on a glass microscope slide and mounted onto an inverted fluorescent microscope (Nikon Eclipse, Ti-E). The pipette tip was aligned with the hydrogel spheres, and basal negative pressure was applied to capture it stably. Aspiration dynamics in response to applied pressure (relative to the basal levels) inside the pipette were recorded (Andor Zyla 4.2 sCMOS) by imaging the Cy3 fluorescent channel using a CFI Super Plan Fluor ELWD 40XC (Nikon). The mechanical properties of the hydrogel spheres were evaluated based on the relationship between the aspirated based length L (t) and the applied pressure ΔP using the half-space model^25^:

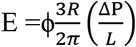

where R is the inner pipette radius and ϕ~2 is the geometrical factor.

### Determination of the size of clones in hydrogel spheres

In order to measure the size of clones developed in PEGDT/MALDEX hydrogel spheres, we took confocal fluorescence microscopy images of clones in hydrogel spheres and counted the number of cells per clone. ESCs and PC9 cells were encapsulated in PEG-DEX hydrogel and grown for 3 to 4 days for ESCs with 2i+LIF and 6 to 7 days for PC9 cells. Spheres containing clones were selected randomly from the tissue culture plate then fixed with 10% v/v formaldehyde (Sigma-Aldrich, cat #F8775). The cells were stained with DAPI Fluoromount-G® (ENCO, cat #0100-20). Cross-sectional images were taken using a NIKON A1 confocal microscope. The numbers of cells per clone were determined by counting the number of nuclei. The surface areas of clones were determined using ImageJ image processing software (NIH, https://imagej.nih.gov/ij/).

### ESC differentiation analyses

For differentiation experiments, ESCs cells were grown in either 2D configuration on 0.1% gelatin-coated standard tissue culture dishes or within hydrogel spheres under indicated conditions. Cells grown in 2D or single cells from hydrogel spheres were washed three times with PBS and resuspended in 1) basal ESC medium supplemented with 1 μM all-*trans* retinoic acid, 2) basal ESC medium without 2i, or 3) basal ESC medium without 2i and LIF for additional 4 days before gene expression profiling evaluation.

### Clonal barcoding and sequencing

To assess the impact of clonal cell origin on cellular states and variation, we produced PC9 cell lines that carry genomic barcodes. We first designed and produced a plasmid pool containing a UGI region of 8 bp as a genomic barcode about 600 bp upstream of the *BFP* polyA region; BFP expression allowed us to detect plasmid integration. The design of plasmid can be found in **Supplementary File**. To validate the number of unique barcodes and their even distribution, we produced a sequencing library from about 50,000 cells by amplifying the plasmid by 12 PCR cycles (2 min at 98 °C, 2 × (98 °C 20 s, 55 °C 30 s, 72 °C 40 s), 10 × (98 °C 20 s, 65 °C 30 s, 72 °C 40 s)). For transfection, we grew 293T cells to 80% confluency and then incubated the cells for 30 min in conditioning medium (50 ml 293T medium supplemented with 500 μl L-glutamine and 500 μl Sodium-Pyruvate). The transfection solution contained 34.5 μl TransIT®-LT1 Transfection Reagent (Mirus, cat #MIR-2300), 1 μg transfer plasmid (psPAX.2), 7 μg VSV-G (PMD2.G), and 3.5 μg of our BFP-UGI plasmid diluted to 1.5 ml with Opti-MEM I Reduced Serum Medium (Gibco, cat #31985088). The transfection solution was incubated for 30 min at room temperature. The transfection solution was then added dropwise onto the cells. Cells were incubated with gentle shaking, and media containing viruses were collected after 48 h and 72 h. For virus concentration, the PEG Virus Precipitation Kit (BioVision, cat #K904) was used. The collected media was PEG precipitated overnight at 4 °C, filtered through a 0.45-μm pore size filter (MF-Millipore), and centrifuged at 2500 rcf for 30 min. The resulting virus pellet was resuspended in 20 μl Virus Resuspension Solution supplied with the kit.

For infection, PC9 cells of 50% confluency were incubated for 30 min in 1 μl polybrene (Sigma-Aldrich, cat #107689 in 1 ml culture medium. The cells were then loaded dropwise with the virus solution and incubated for 2.5 h in a humidified incubator at 37 °C, 5% CO2. PC9 cells were infected with the virus at an MOI of 1. Finally, BFP-positive cells were sorted by FACS to obtain the BFP PC9 cell line containing genetic barcodes. We encapsulated BFP PC9 cells in PEGDT/MALDEX hydrogel and grew them for 7 days. Hydrogel spheres were washed and degraded as described above, and the released cells were trypsinized and resuspended to obtain a single-cell solution, which was subjected to scRNA-seq.

### Single-cell/clone microfluidic droplet barcoding using inDrops

Single cells and single clone transcriptomes were barcoded using inDrops as previously reported^6^. **Device III** was used for scRNA-seq, and **device IV** was used for CloneSeq (**Supplementary Figure 3C-D**). The devices have four inlets: **1) Cell/encapsulated clone inlet**: *For **scRNA-seq experiments***, cells were loaded at 200,000 cells/ml in PBS containing 10% v/v OptiPrep and maintained in suspension using a magnetic micro-stirrer bar placed within the tube. ***For CloneSeq experiments***, around 20,000 clones were resuspended in 900 μl PBS and 100 μl OptiPrep and maintained in suspension using a magnetic micro-stirrer bar placed within the tube. **2) Barcoding acrylamide beads inlet**: Barcoding beads were prepared as previously described^29^ and kept in dark at 4 °C in 50% (v/v) 10 mM Tris-HCl (pH 8), 0.1 M EDTA, 0.1% (v/v) Tween-20. Around 100-150 μl barcoded acrylamide beads were centrifuged in a 1.5-ml Eppendorf tube at 1500 rcf for 2 min to obtain packed beads. After aspirating residual buffer from the pelleted beads, the tube was loaded onto the corresponding inlet in the microfluidics setup. **3) Reverse transcription/lysis mix inlet:***For both single-cell and CloneSeq* **experiments**, the RT/lysis mix consisted of 180 μL 5X First-Strand buffer (SuperScript™ III Reverse Transcriptase Kit, Invitrogen Cat #18080044), 27 μL 10% (v/v) IGEPAL CA-630 (Sigma-Aldrich, Cat #I8896), 20 μL 25 mM dNTPs (NEB, Cat #N0446S), 30 μL 0.1 M DTT (SuperScript™ III Reverse Transcriptase Kit, Invitrogen Cat #18080044), 45 μL 1 M Tris-HCl (pH 8.0) (Sigma-Aldrich, Cat #T2319), 30 μL murine RNase inhibitor (NEB, Cat #M0314), 45 μL SuperScript™ III RT enzyme (200 U/μL, Invitrogen Cat #18080044), and 73 μL nuclease-free water (Sigma-Aldrich, Cat #W4502). **4) Carrier oil inlet:** The carrier oil was 3 ml of HFE-7500 with 2% (w/w) fluorosurfactant. During microfluidics runs, cell suspension/encapsulated cells and collection tubes were kept on ice. The device generates monodispersed droplets with volumes in the range of 2 nl for scRNA-seq and around 4-5 nl for CloneSeq. The flow rates used for sequencing are shown in **Supplementary Table 1**.

### Single-cell/clone microfluidic droplet barcoding using Drop-seq

Single cells and single clone transcriptomes were barcoded using Drop-seq as previously reported^30^. We used the same devices used for InDrops (**Device III** for scRNA-seq; **device IV** for CloneSeq) while plugging port No. 7 (**Supplementary Figure 3C-D**). The devices have three inlets: **1) Cell/clone inlet**: Cells were loaded at 12,000 cells/ml, and clones were loaded at 5,000 clones/ml in PBS containing 10% v/v OptiPrep and were maintained in suspension using a magnetic micro-stirrer bar placed within the tube. **2) Barcoding/lysis mix inlet**: An aliquot of barcode beads (Chemgenes Corp.) containing 300,000 barcodes at a concentration of ~400 beads/μL was removed from the stock tube and washed twice with 1 ml lysis solution made of 67.5 μL 10% (v/v) IGEPAL CA-630, 112.5 μL 1 M Tris-HCl (pH 8.0), and 820 μL nuclease-free water. The beads were then resuspended in 1.5 ml 100 μL 10% (v/v) IGEPAL CA-630, 112 μL 0.1 M DTT, 170 μL 1 M Tris-HCl (pH 8.0), 118 μL murine RNase inhibitor, and 1 ml nuclease-free water leading to a concentration of 200,000 barcode beads/ml. Following resuspension, the sample was loaded onto the corresponding inlet in the microfluidics setup. **3) Carrier oil inlet:** The carrier oil was 10 ml of HFE-7500 with 2% (w/w) fluorosurfactant. During microfluidics runs, cell suspension/encapsulated cells, lysis mix, and collection tubes were kept on ice. The device generates monodispersed droplets with volumes in the range of 0.5 nl for scRNA-seq and 1 nl for CloneSeq. Flow rates used for Drop-seq are shown in *Supplementary Table 1.*

### Library preparation

The Drop-seq libraries were prepared following a previously published protocol30. All the primers used in the library preparation are listed in Supplementary Table 2. The inDrops libraries were prepared using the following procedure: After completion of the microfluidics stage, the collection tubes were exposed to 6.5 J/cm2 of a 365-nm UV lamp for 10 min to release photocleavable barcoding primers from the barcoding beads. Next, the collection tubes containing the UV-exposed emulsion were transferred to a reverse transcription reaction at 50 °C for 2 h followed by 15 min at 70 °C to stop the reaction. Each sample was then demulsified by adding 50 μl PFO to release the barcoded cDNA from the droplets. After clear separation of the two phases was observed, the upper aqueous phase containing the barcoded cDNA was transferred to a new well. To remove unused primers and primer dimers, a 1:1 digestion mix was added, containing 20 U/μL Exonuclease I (NEB, Cat #M0293), 20 U/μL HinfI enzyme (NEB, Cat #R0155), x1 Exonuclease I Reaction Buffer (NEB, Cat #B0293), x1 CutSmart buffer (NEB, Cat #B7204), and 30 μl of nuclease-free water. Samples were incubated for 1 h at 37 °C and 10 min at 80 °C. The reaction product (in the form of a cDNA:RNA hybrid) was purified with a 1.5X reaction volume of AMPure XP beads (Beckman Coulter, Cat #A63882) and eluted in 13.5 μl TE buffer. For second strand synthesis, 13.5 μl digestion reaction product was combined with 1.5 μl second-strand synthesis (SSS) buffer and 1 μl of SSS enzyme mix from the NEBNext mRNA Second Strand Synthesis Module (NEB, Cat #E6111) and incubated at 16 °C for 2.5 h, followed by 20 min at 65 °C. For linear amplification by *in vitro* transcription, SSS reaction products (16 μl) were combined with 24 μl T7 High Yield RNA Synthesis Kit (NEB, Cat #E2040) reagent mix containing 4 μl T7 Buffer, 4 μl ATP, 4 μl CTP, 4 μl GTP, 4 μl UTP, and 4 μl T7 enzyme mix. The reaction was incubated at 37 °C for 13 h, and the resulting RNA was purified with 1.3x reaction volume of AMPure XP beads and eluted with 20 μl TE buffer. An aliquot of 9 μl was frozen for backup at −80 °C, a 2-μl sample was taken for direct analysis, and the remaining 9 μl was used in subsequent library preparation steps. Next, RNA was fragmented using an RNA fragmentation kit (Invitrogen, Cat #AM8740). The 9-μl aliquot of RNA were combined with 1 μl of RNA fragmentation reagent and incubated at 70 °C for 2 min, transferred to ice, and 40 μl fragmentation stop mix containing 5 μl fragmentation stop solution and 35 μl TE buffer was added. Fragmented RNA was purified with a 1.3X reaction volume of AMPure XP beads and eluted in 10 μl TE buffer. The resulting amplified and fragmented RNA was reverse transcribed using a random hexamer primer as follows: first, 10 μl RNA was mixed with 2 μl of 100 μM PE2-N6-v2 random hexamer primer (5’-AGACGTGTGCTCTTCCGATCTNNNNNN-3’) and 1 μl of 10 mM dNTPs, incubated for 3 min at 65 °C and transferred to ice. Then the following components were added to the reaction: 4 μl of 5X First-Strand buffer, 1 μl of 0.1 M DTT, 1 μl murine RNase inhibitor, and 1 μl of SuperScript™ III RT enzyme (200 U/μL). Samples were incubated at 25 °C for 5 min, 50 °C for 60 min, and 70 °C for 15 min. For the clonal barcoding library, UGI-shifted primers (5’-AGACGTGTGCTCTTCCGATCTGTCGACGGATCC-3’) were used instead of random primers for half of the sample to amplify the genetic barcode region with incubation at 48 °C for 5 min, 55 °C for 60 min, and 70 °C for 15 min. Following reverse transcription, the reaction volume was raised to 50 μl by adding 30 μl nuclease-free water, and the resulting cDNA was purified with 1.2X reaction volume of AMPure XP beads and eluted in 11.5 μl TE buffer. The resulting libraries were PCR amplified using standard PE1/PE2 full-length primer mix (2p fixed: 5’-AATGATACGGCGACCACCGAGATCTACACTCTTTCCCTACACGACGCTCTTCCGATCT-3’ 2p fixed + barcode: 5’-CAAGCAGAAGACGGCATACGAGATNNNNNNNNGTGACTGGAGTTCAGACGTGTGCTCTT CCGATCT-3’). The primers contain Illumina library indices for multiplexing. Each PCR reaction consisted of 14 amplification cycles and contained 11.5 μl post-reverse transcription cDNA library, 12.5 μl 2x KAPA HiFi HotStart ReadyMix (Roche), and 1 μl of 25 μM PE1/PE2 index primer mix. Amplified libraries were purified using a 0.7X reaction volume of AMPure XP beads and eluted in 30 μl nuclease-free water. Aliquots of 15 μl of each resulting library were run in 2% agarose gels, and the desired 200-800 bp DNA library fragments were isolated using PureLink™ Quick Gel Extraction Kit (Invitrogen, cat #K210012). For the clonal barcoding library, we aimed for amplification of a band around 600 bp. Library quality was confirmed by Agilent 2200 TapeStation nucleic acid system (Agilent) using the Agilent High Sensitivity D1000 DS DNA kit. The resulting libraries had an average size of 350-550 bp (Supplementary Figure 7). Size-selected libraries were diluted to 4 nM and combined into a pool for paired-end, single index sequencing on the Illumina NextSeq 550 instrument, using an Illumina 550 High Output v2 (75 cycles) kit. Cycle distribution was 45 cycles for Read 1, 35 cycles for Read 2, and 8 cycles for library index read.

**Supplementary Table 2.**
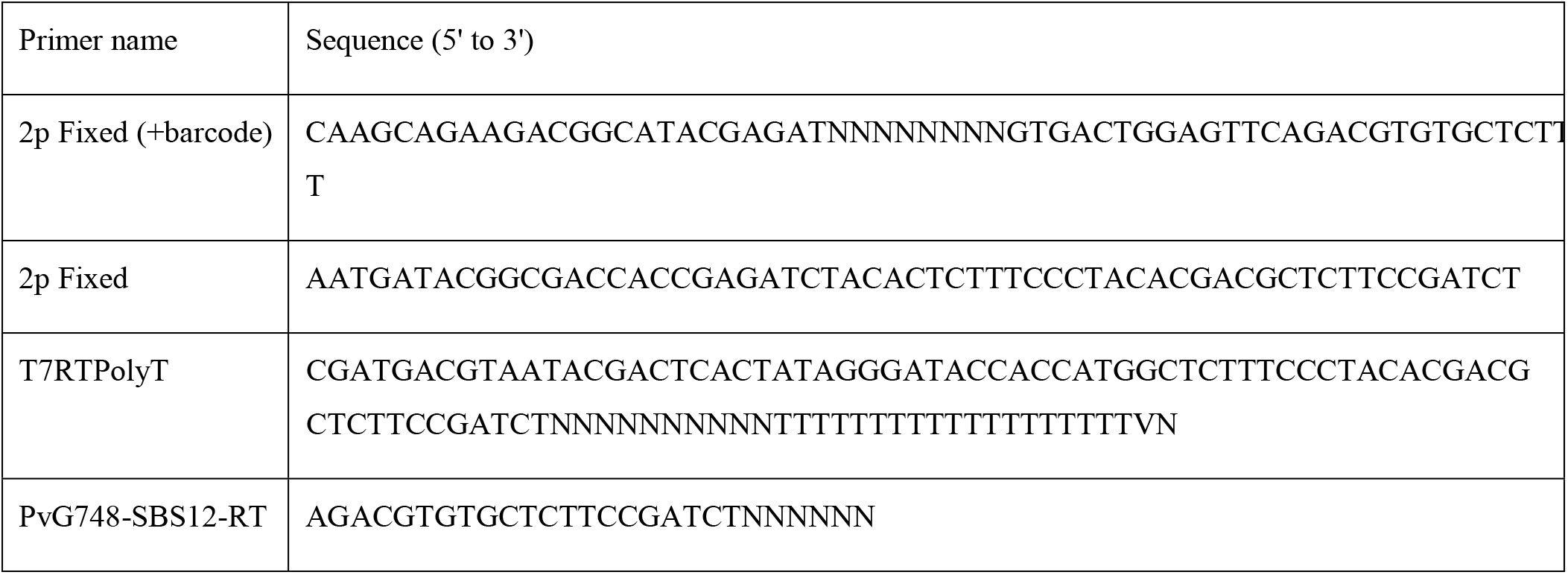
Primers used in library preparation.

### Species-mixing experiments

To determine off-species contamination in our single-cell and clonal preparations, we performed inDrops as described above with a PC9/R1 ESCs cell suspension mixture. The suspension mixtures were 100,000 cells/ml in total (1:1 human:mouse ratio) for the single-cell experiment and 20,000 clones/ml in total (1:1 ratio) for single clone experiment. PC9 cells were identified as those barcodes with greater than 15,000 human transcripts, and R1 mESCs were identified as those with greater than 15,000 mouse transcripts.

### Sequencing and data filtering

Paired-end sequencing was performed on Illumina NextSeq 500. Read 1 was used to obtain the sample barcode and UMI sequences, and read 2 was mapped to a reference transcriptome. The reads were first filtered based on the presence of two sample barcode components separated by the W1 adaptor sequence in Read 1. Barcodes for each read were matched against a list of the 3842 pre-determined barcodes, and errors of up to two nucleotides mismatch were corrected. Reads with a barcode separated by more than two nucleotides from the reference list were discarded. The reads were then split into barcode-specific files for mapping and UMI filtering.

### Clonal barcoding analysis

For clonal barcoding experiments, two libraries were generated from the same sample: One was a general library made using a random primer that showed the transcription profile background, whereas the second was made using the UGI-shifted primer that only presented a narrow region around the clone barcodes. The list was matched against the pre-determined barcode reference list, and errors of up to two nucleotides were corrected. Reads with a barcode separated by more than two nucleotides from the reference list were discarded. Some cells carried more than one clone barcode as a result of an MOI greater than 1 at the infection step. Clonal cell origin was determined by matching cell UMIs from the random library and the UGI library. To compare similarities between clonal cells to cells picked at random from the whole population, we first created a tSNE plot of the random primer library as background, then marked the clonal origins of cells on the plot. The Euclidian distance between cells was calculated and compared between clonal cells and random cells first by coordinates on tSNE plot then by the entire gene expression matrix. The significance of the difference between the distance within clones and random cells was tested by a Wilcoxon rank sum test.

### Single clone alignment and UMI-based filtering

Reads split into clone barcode-specific files were aligned using Bowtie2 to the mouse or human reference transcriptome. Alignments from Bowtie were filtered as follows: (1) For each read, we retained at most one alignment per gene, across all isoforms, by choosing the alignment closest to the end of the transcript. (2) If a read aligned to multiple genes, we excluded any alignments more than 400 bp away from the end of the transcript. This step results in an approximately 5% increase in the number of final UMI reads obtained, as compared to simply discarding any ambiguous read. (3) If a read still aligned to more than two genes after UMI filtering, we excluded the read altogether.

### Gene set signature activation analysis

We used MSigDB GSEA mapped to NCI-60 cell lines using GO biological processes and REACTOME gene sets with FDR q-value less than 0.01 as described previoulsy^38,107^.

### Deep sequencing

Deep sequencing was carried out on an Illumina NextSeq using commercially available kits from Illumina (Danyel Biotech, Cat #FC-404-2005) following the manufacturer’s protocols.

### Data analysis

The Illumina output was analyzed using an in-house Perl script that produced a reads matrix that was aligned using RSEM ^108^ with Bowtie ^109^. The resulting matrix was analyzed in R. For bulk data analysis the transcript per million (TPM) values were used to compare libraries. Differential gene expression was visualized using xy plots. Statistical analysis was performed for replicates using a two-sided t-test, and *p* values of <0.05 were deemed significant.

scRNA-seq data was analyzed using the Seurat v2.4 pipeline ^35^. For single cells, cells with more than 5,000 unique molecular identifiers were retained for further analysis. Clones with more than 15,000 unique molecular identifiers were retained for further analysis. A global-scaling normalization was performed on the filtered dataset using “LogNormalize” with a scale factor of 10,000. Identification of highly variable genes was performed with the following parameters: x.low.cutoff = 0.2, x.high.cutoff = 5, y.cutoff = 0.5, and y.high.cutoff = 10. Cell-to-cell variation in gene expression driven by batch, cell alignment rate, and the number of detected molecules were regressed out and a linear transformation was applied. A principal component analysis was performed on the scaled data with 15 principal components. Identification of clusters of cells was done by a SNN modularity optimization based clustering algorithm. We first calculated *k*-nearest neighbors and then constructed the SNN graph. The modularity function was optimized to identify clusters. Clustering was done with resolution of 0.6, and tSNE or UMAP was used for visualization.

---

**Supplementary Figure 1.**
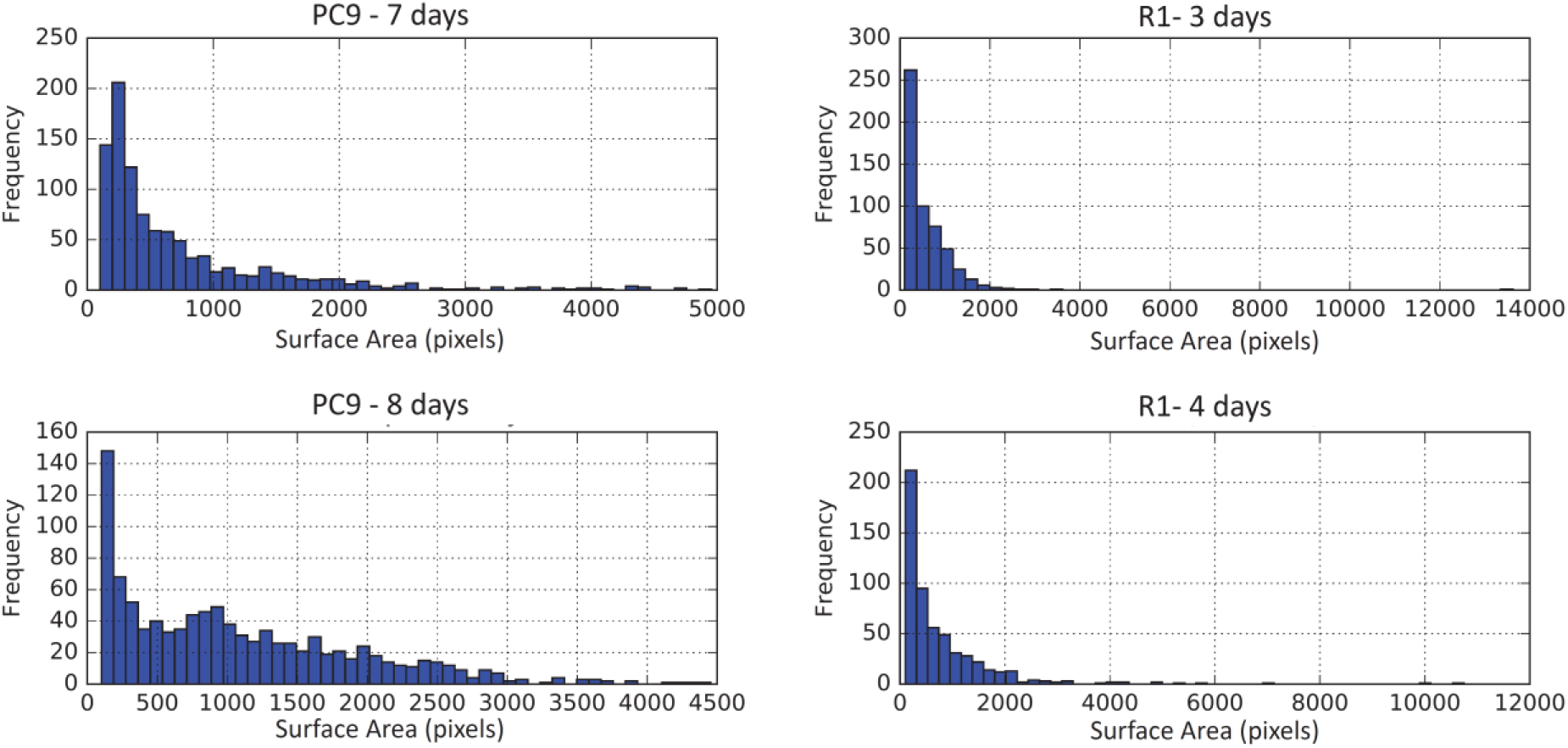
Distribution of clone sizes. The size distributions of clones based on surface area in microfluid photos showing a strong bias towards smaller sizes. For PC9, clones were grown for 7 and 8 days. For ESCs, clones were grown for 3 and 4 days.

**Supplementary Figure 2.**
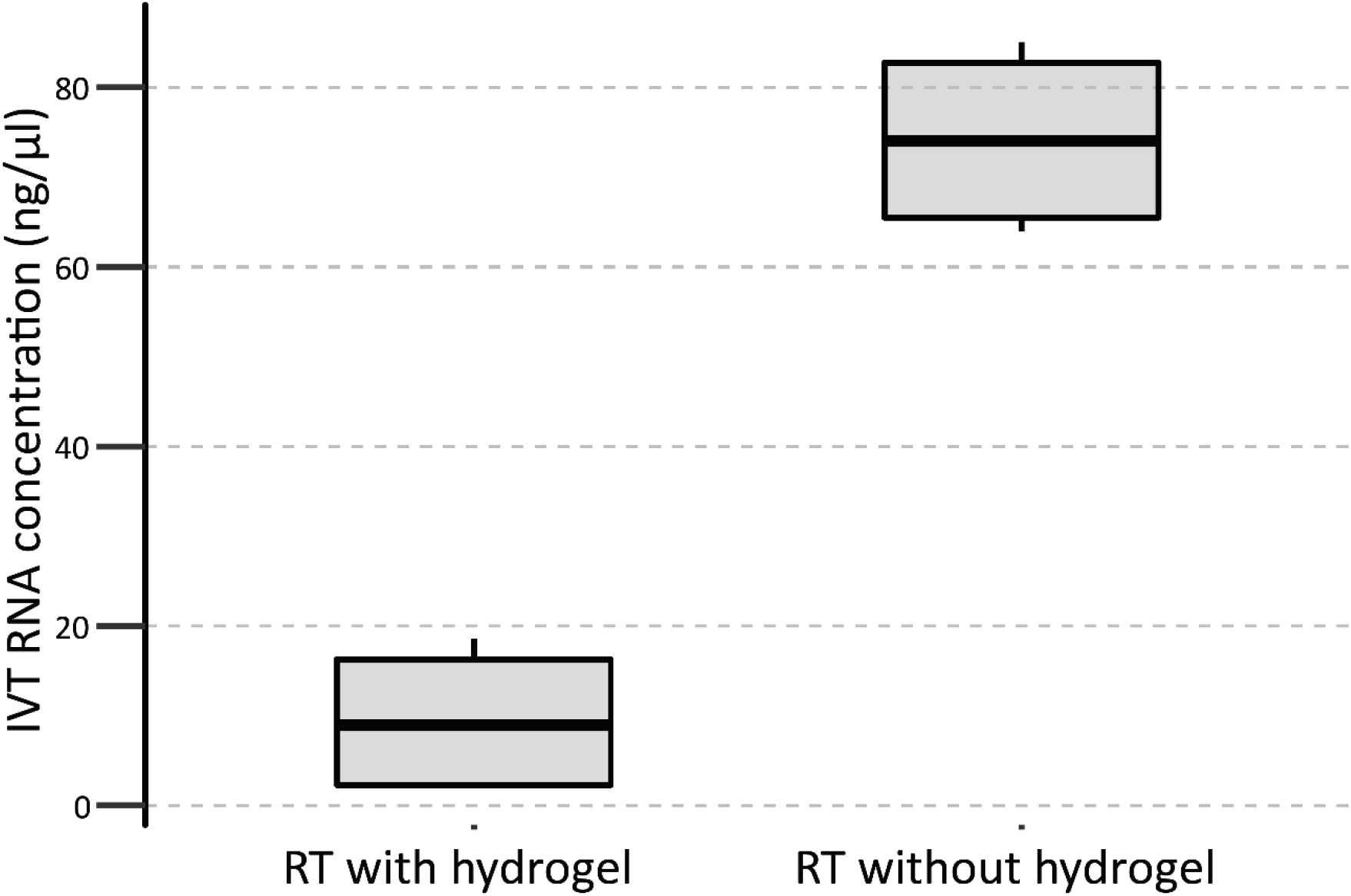
Inhibition of reverse transcriptase by the presence of PEGDT/MALDEX hydrogel. The efficiency of reverse transcription reaction of total RNA extracted from bulk PC9 cells in the presence and absence of empty PEGDT/MALDEX hydrogel, as determined by the concentration of RNA synthesized in the in-vitro amplification step.

**Supplementary Figure 3.**
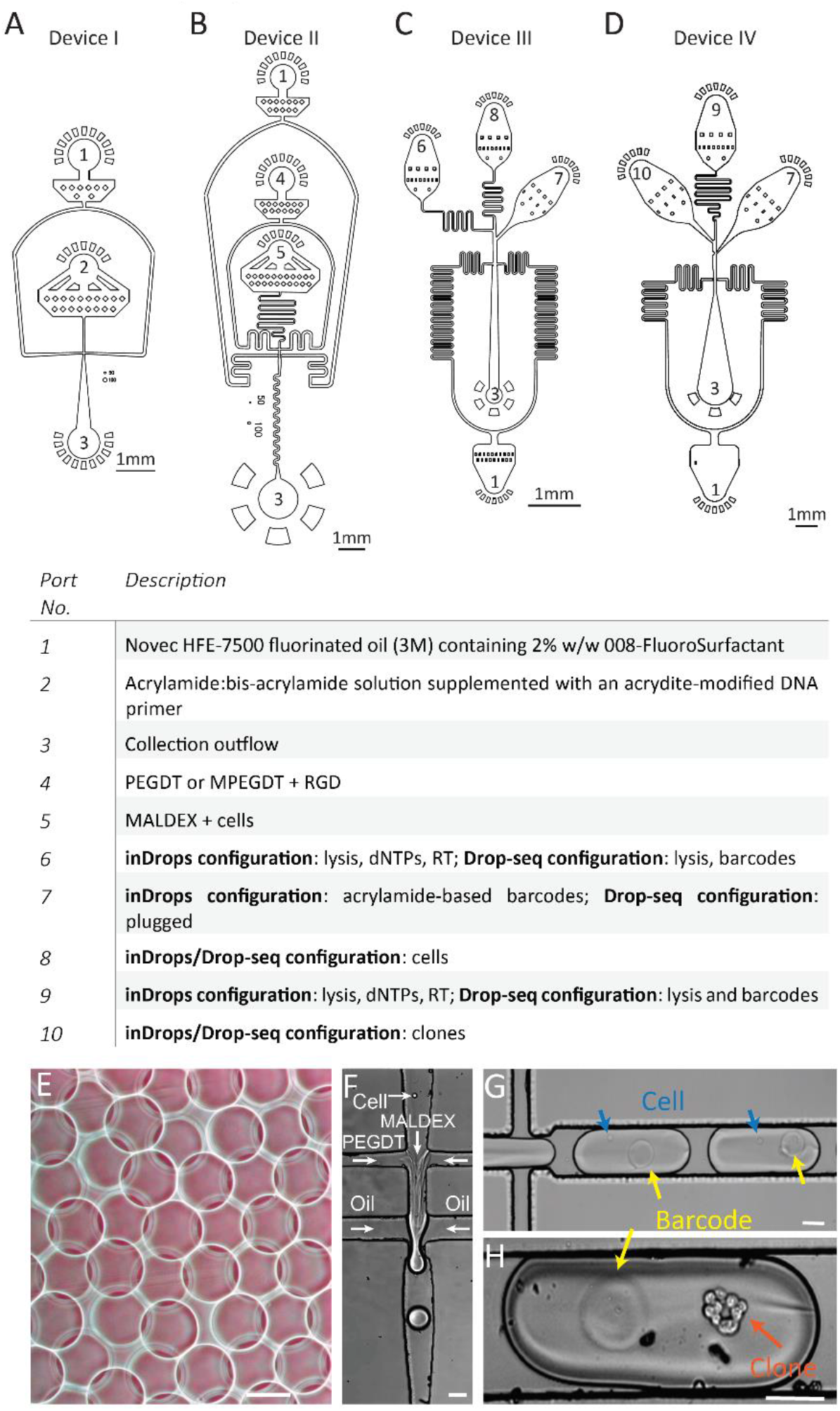
Microfluidic devices used for (**A**) acrylamide-based barcode fabrication, (**B**) cell encapsulation within PEGDT/MALDEX hydrogel; (**C**) scRNA-seq, and (**D**) single-clone RNA seq. (**E**) Barcode beads produced by device I. (**F**) Image of a single-cell encapsulation within PEGDT/MALDEX hydrogel using device II. (**G**) Image of single cells captured with drops using device III. (**H**) Image of clone captured within a drop using device IV. Scale bars: e-h: 50 μm.

**Supplementary Figure 4.**
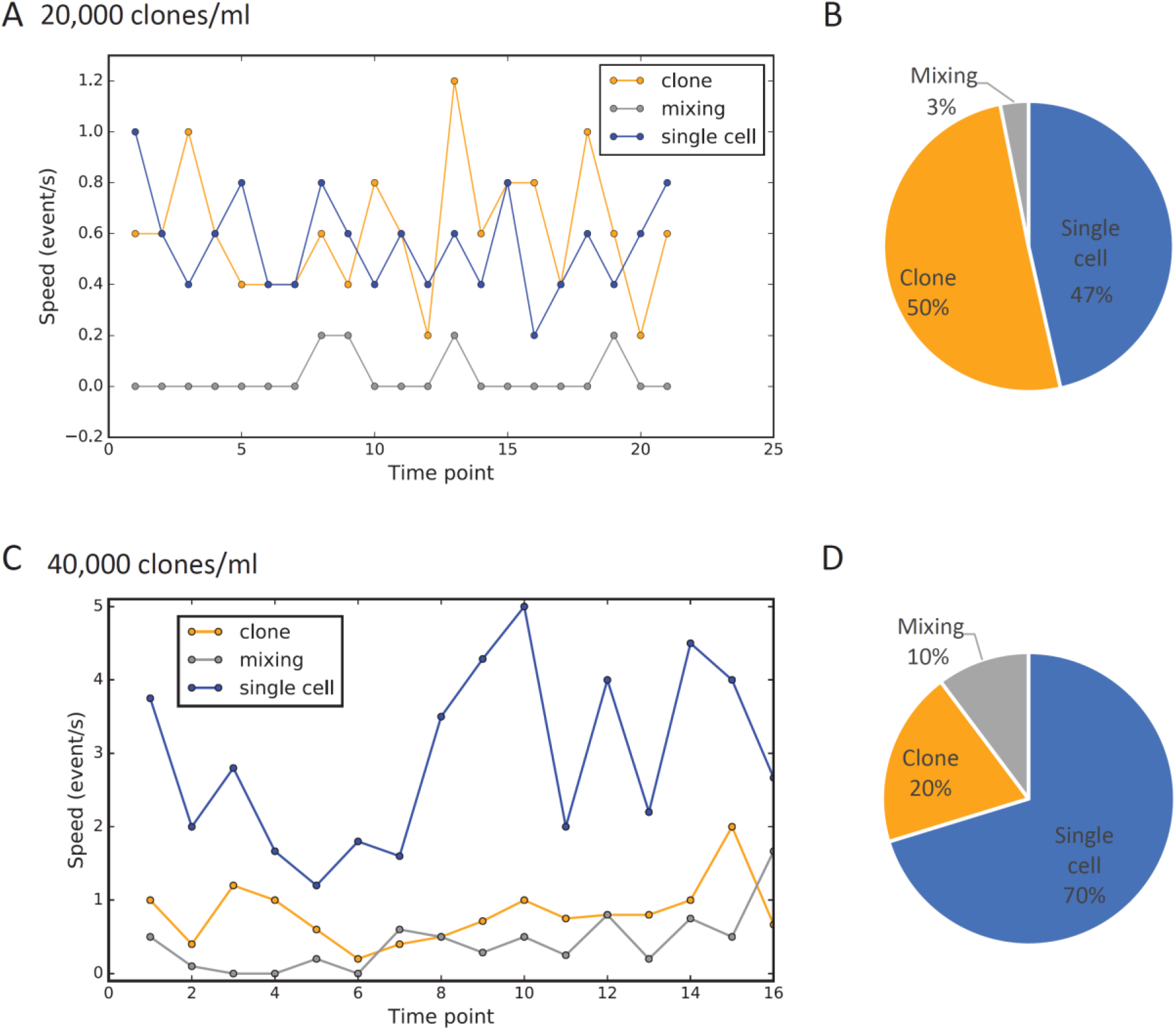
Counting of mixing events during CloneSeq encapsulation. Counting was based on in-time microscope videos at the encapsulation junction of the device during CloneSeq. Each time point in axis-x (**A and C**) was calculated by counting the number of events per five seconds for capturing a single cell, a single clone or a mixed event. Two concentrations were tested. 20,000 clones/ml (**A-B**) and 40,000 clone/ml (**C-D**). Pie charts summarize the counts in each concentration tested (**B and D**). Overall, mixing rate was as low as 3% with 20,000 clones/ml (**B**) and 10% with 40,000 clones/ml (**D**).

**Supplementary Figure 5.**
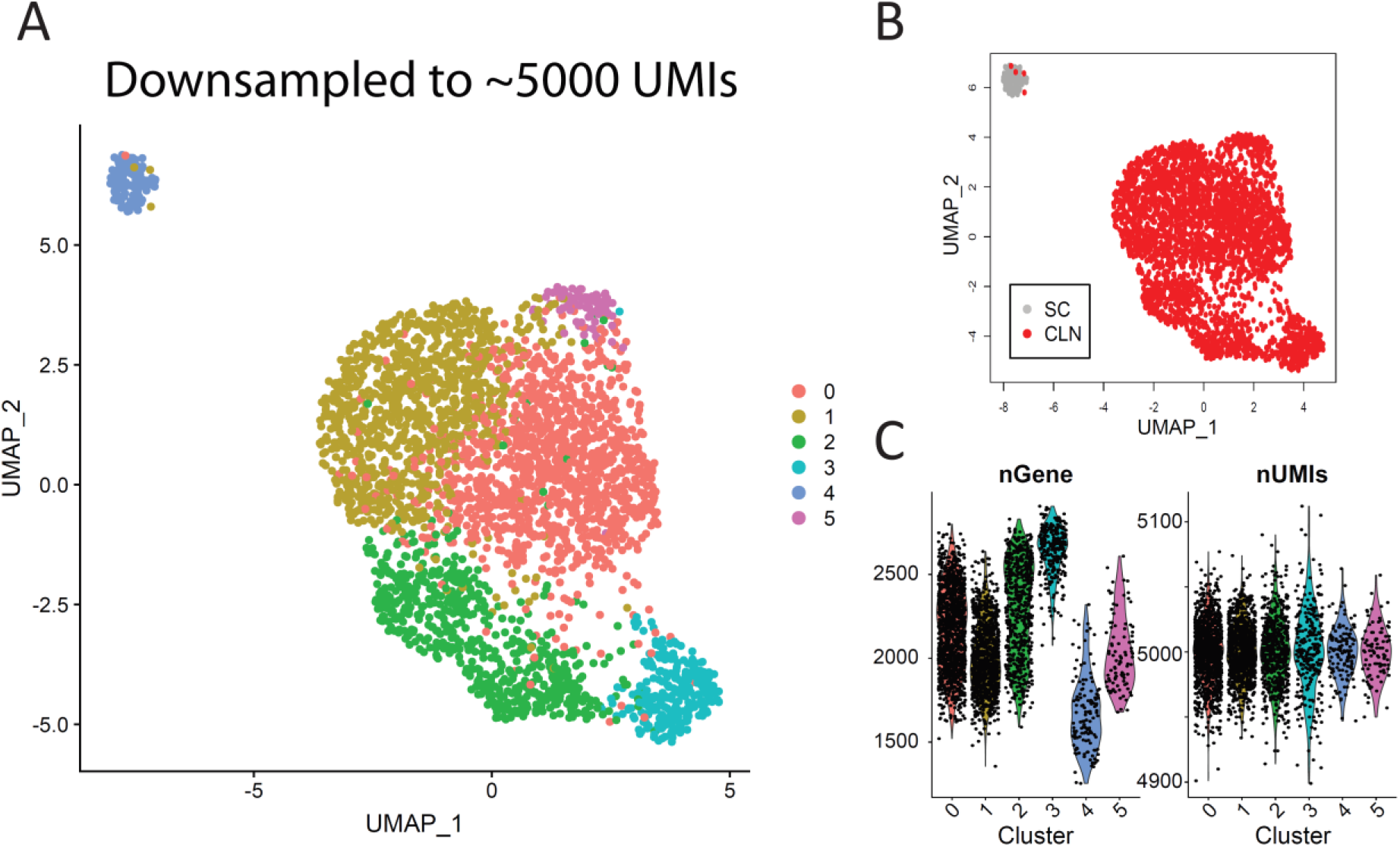
The plot shows UMAP for 3000 clones and 300 single cells, same as figure 4, after UMI reduction by downsampling each clone and single cell to 5,000 UMIs. (**A**) UMAP colored by cluster. (**B**) UMAP colored based on clones vs single cells (CLN = CloneSeq, SC = scRNA-seq). (**D**) Violin plots of distributions of the numbers of genes (left) and transcripts (scored by UMIs) divided into clusters.

**Supplementary Figure 6.**
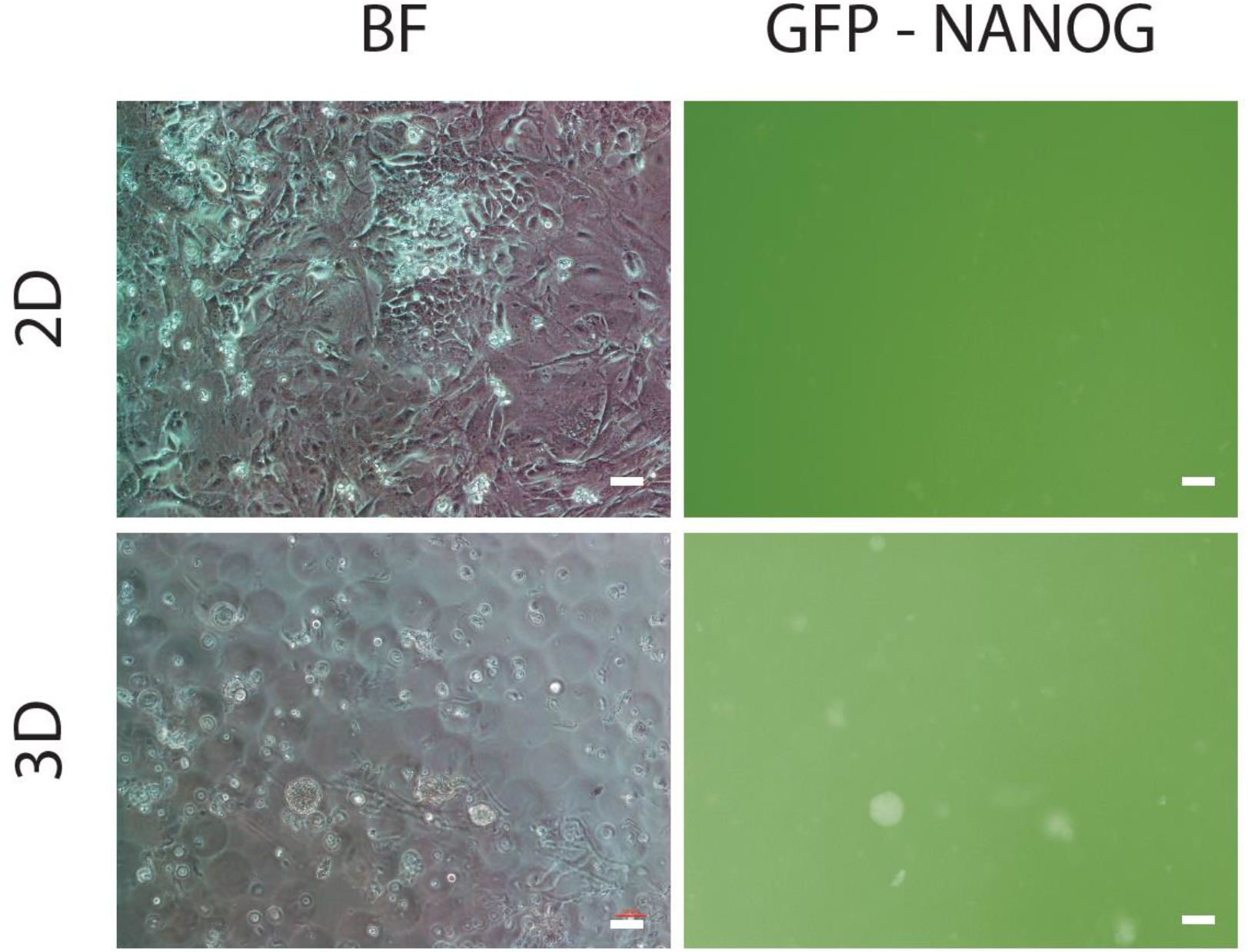
MEFs containing the OSKM cassette under the control of the TET-on promoter were induced into pluripotent stem cells (iPSCs) in spheres (3D) and on gelatin (2D). Images show cells after 4 days of Dox inducing reprogramming. Scale bars: 50μm.

**Supplementary Figure 7.**
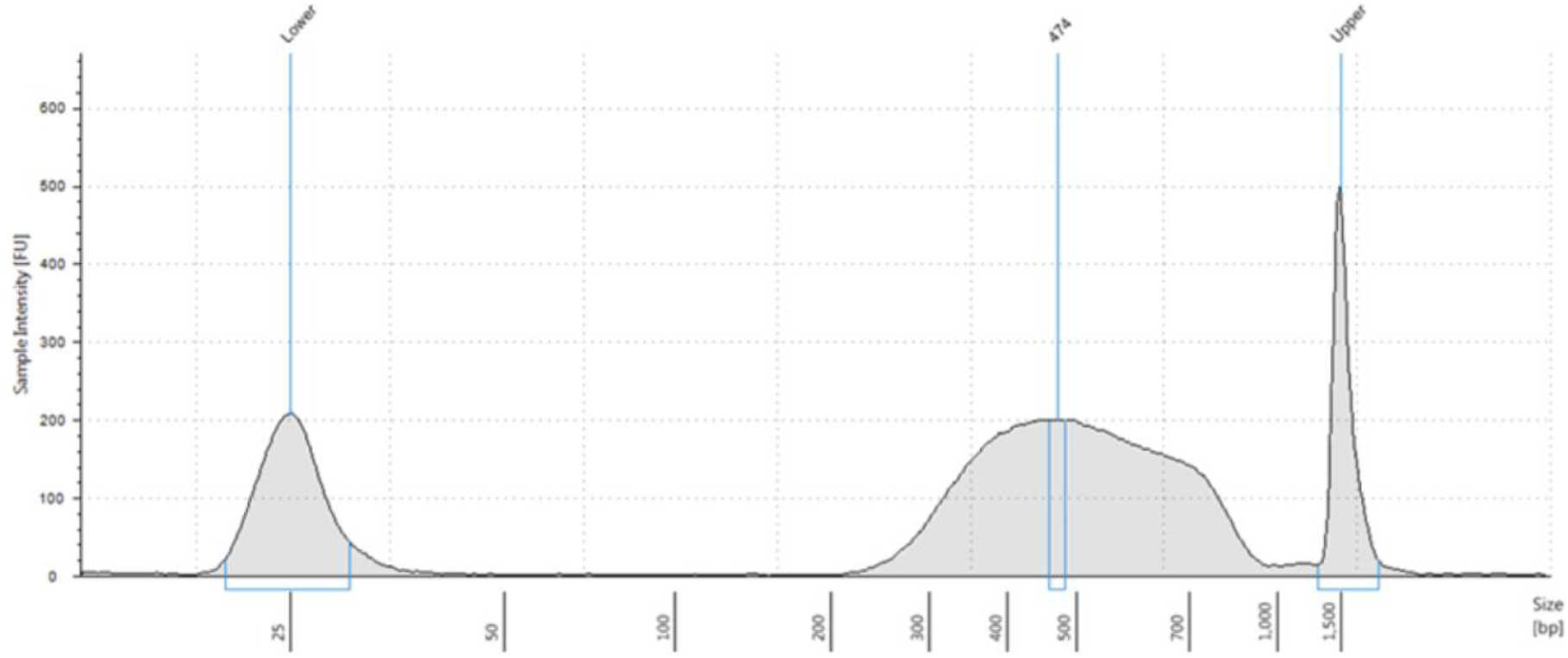
Representative Agilent by 2200 TapeStation electropherogram of a typical ESCs CloneSeq library after size selection on 2% agarose gel.

## Notes

### Competing Interest Statement

The authors have declared no competing interest.

## References

1. Kim, K.-T. et al. Single-cell mRNA sequencing identifies subclonal heterogeneity in anti-cancer drug responses of lung adenocarcinoma cells. Genome Biol. 16, 127 (2015).

2. Dalerba, P. et al. Single-cell dissection of transcriptional heterogeneity in human colon tumors. Nat. Biotechnol. 29, 1120–1127 (2011).

3. Li, H. et al. Reference component analysis of single-cell transcriptomes elucidates cellular heterogeneity in human colorectal tumors. Nat. Genet. 49, 708–718 (2017).

4. Tirosh, I. et al. Dissecting the multicellular ecosystem of metastatic melanoma by single-cell RNA-seq. Science (80-.). 352, (2016).

5. Yan, L. et al. Single-cell RNA-Seq profiling of human preimplantation embryos and embryonic stem cells. Nat. Struct. Mol. Biol. 20, 1131–1139 (2013).

6. Klein, A. M. et al. Droplet barcoding for single-cell transcriptomics applied to embryonic stem cells. Cell 161, 1187–1201 (2015).

7. Patel, A. P. et al. Single-cell RNA-seq highlights intratumoral heterogeneity in primary glioblastoma. Science (80-.). 344, 1396–1401 (2014).

8. Ma, K.-Y. Y. et al. Single-cell RNA sequencing of lung adenocarcinoma reveals heterogeneity of immune response–related genes. JCI insight 4, (2019).

9. Chung, W. et al. Single-cell RNA-seq enables comprehensive tumour and immune cell profiling in primary breast cancer. Nat. Commun. 8, 1–12 (2017).

10. Martello, G. & Smith, A. The Nature of Embryonic Stem Cells. Annu. Rev. Cell Dev. Biol. (2014) doi:10.1146/annurev-cellbio-100913-013116.

11. Evans, M. J. & Kaufman, M. H. Establishment in culture of pluripotential cells from mouse embryos. Nature 292, 154–156 (1981).

12. Raj, A. & van Oudenaarden, A. Single-Molecule Approaches to Stochastic Gene Expression. Annu. Rev. Biophys. 38, 255–270 (2009).

13. Chen, S., Lake, B. B. & Zhang, K. High-throughput sequencing of the transcriptome and chromatin accessibility in the same cell. Nature Biotechnology vol. 37 1452–1457 (2019).

14. Litzenburger, U. M. et al. Single-cell epigenomic variability reveals functional cancer heterogeneity. Genome Biol. 18, 15 (2017).

15. Pan, X. N. et al. Inhibition of c-Myc overcomes cytotoxic drug resistance in acute myeloid leukemia cells by promoting differentiation. PLoS One 9, (2014).

16. Losick, R. & Desplan, C. Stochasticity and cell fate. Science vol. 320 65–68 (2008).

17. Dar, R. D. et al. Transcriptional burst frequency and burst size are equally modulated across the human genome. Proc. Natl. Acad. Sci. U. S. A. (2012) doi:10.1073/pnas.1213530109.

18. Nitzan, M., Karaiskos, N., Friedman, N. & Rajewsky, N. Gene expression cartography. Nature 576, 132–137 (2019).

19. Tung, P. Y. et al. Batch effects and the effective design of single-cell gene expression studies. Sci. Rep. 7, (2017).

20. Ziegenhain, C. et al. Comparative Analysis of Single-Cell RNA Sequencing Methods. Mol. Cell 65, 631–643.e4 (2017).

21. Nuttelman, C. R., Mortisen, D. J., Henry, S. M. & Anseth, K. S. Attachment of fibronectin to poly(vinyl alcohol) hydrogels promotes NIH3T3 cell adhesion, proliferation, and migration. J. Biomed. Mater. Res. 57, 217–23 (2001).

22. Li, C. Y. et al. Micropatterned cell-cell interactions enable functional encapsulation of primary hepatocytes in hydrogel microtissues. Tissue Eng. Part A 20, 2200–12 (2014).

23. Diamandis, E. P. & Christopoulos, T. K. The biotin-(strept)avidin system: principles and applications in biotechnology. Clin. Chem. 37, 625–36 (1991).

24. Hochmuth, R. M. Micropipette aspiration of living cells. Journal of Biomechanics vol. 33 15–22 (2000).

25. Theret, D. P., Levesque, M. J., Sato, M., Nerem, R. M. & Wheeler, L. T. The application of a homogeneous half-space model in the analysis of endothelial cell micropipette measurements. J. Biomech. Eng. (1988) doi:10.1115/1.3108430.

26. Bainer, R. & Weaver, V. Strength under tension. Science vol. 341 965–966 (2013).

27. Burdick, J. A. & Murphy, W. L. Moving from static to dynamic complexity in hydrogel design. Nat. Commun. 3, 1269 (2012).

28. Almalki, S. G. & Agrawal, D. K. Effects of matrix metalloproteinases on the fate of mesenchymal stem cells. Stem Cell Res. Ther. 7, 129 (2016).

29. Zilionis, R. et al. Single-cell barcoding and sequencing using droplet microfluidics. Nat. Protoc. 12, 44–73 (2017).

30. Macosko, E. Z. et al. Highly Parallel Genome-wide Expression Profiling of Individual Cells Using Nanoliter Droplets. Cell 161, 1202–1214 (2015).

31. Rotem, A. et al. Single-cell ChIP-seq reveals cell subpopulations defined by chromatin state. Nat. Biotechnol. 33, 1165–1172 (2015).

32. Rotem, A. et al. High-Throughput Single-Cell Labeling (Hi-SCL) for RNA-Seq Using Drop-Based Microfluidics. PLoS One 10, e0116328 (2015).

33. Tapias, L. F. et al. Assessment of Proliferation and Cytotoxicity in a Biomimetic Three-Dimensional Model of Lung Cancer. Ann. Thorac. Surg. (2015) doi:10.1016/j.athoracsur.2015.04.035.

34. Ma, I. & Allan, A. L. The Role of Human Aldehyde Dehydrogenase in Normal and Cancer Stem Cells. Stem Cell Reviews and Reports (2011) doi:10.1007/s12015-010-9208-4.

35. Stuart, T. et al. Comprehensive Integration of Single-Cell Data. Cell 177, 1888–1902.e21 (2019).

36. Butler, A., Hoffman, P., Smibert, P., Papalexi, E. & Satija, R. Integrating single-cell transcriptomic data across different conditions, technologies, and species. Nat. Biotechnol. 36, 411–420 (2018).

37. Piñero, J. et al. The DisGeNET knowledge platform for disease genomics: 2019 update. Nucleic Acids Res. (2020) doi:10.1093/nar/gkz1021.

38. Subramanian, A. et al. Gene set enrichment analysis: A knowledge-based approach for interpreting genome-wide expression profiles. Proc. Natl. Acad. Sci. U. S. A. (2005) doi:10.1073/pnas.0506580102.

39. Mohammadinejad, R. et al. EMT signaling: potential contribution of CRISPR/Cas gene editing. Cellular and Molecular Life Sciences (2020) doi:10.1007/s00018-020-03449-3.

40. Pradella, D., Naro, C., Sette, C. & Ghigna, C. EMT and stemness: Flexible processes tuned by alternative splicing in development and cancer progression. Molecular Cancer (2017) doi:10.1186/s12943-016-0579-2.

41. Choi, E. J. et al. FOXP1 functions as an oncogene in promoting cancer stem cell-like characteristics in ovarian cancer cells. Oncotarget (2016) doi:10.18632/oncotarget.6510.

42. Edupuganti, R. R. et al. Alternative SET/TAFI Promoters Regulate Embryonic Stem Cell Differentiation. Stem Cell Reports (2017) doi:10.1016/j.stemcr.2017.08.021.

43. Campos, B. et al. Differentiation therapy exerts antitumor effects on stem-like glioma cells. Clin. Cancer Res. 16, 2715–2728 (2010).

44. Mallm, J. P. et al. Glioblastoma initiating cells are sensitive to histone demethylase inhibition due to epigenetic deregulation. Int. J. Cancer 146, 1281–1292 (2020).

45. Ben-David, U. et al. Elimination of undifferentiated cancer cells by pluripotent stem cell inhibitors. J. Mol. Cell Biol. 6, 267–269 (2014).

46. Herreros-Pomares, A. et al. Lung tumorspheres reveal cancer stem cell-like properties and a score with prognostic impact in resected non-small-cell lung cancer. Cell Death Dis. 10, 1–14 (2019).

47. Yu, X., Zhang, Y., Wu, B., Kurie, J. M. & Pertsemlidis, A. The miR-195 Axis Regulates Chemoresistance through TUBB and Lung Cancer Progression through BIRC5. Mol. Ther. - Oncolytics (2019) doi:10.1016/j.omto.2019.07.004.

48. Zeng, L., O’Connor, C., Zhang, J., Kaplan, A. M. & Cohen, D. A. IL-10 promotes resistance to apoptosis and metastatic potential in lung tumor cell lines. Cytokine (2010) doi:10.1016/j.cyto.2009.11.015.

49. Jeon, S. J. et al. TIPRL potentiates survival of lung cancer by inducing autophagy through the eIF2α-ATF4 pathway. Cell Death Dis. (2019) doi:10.1038/s41419-019-2190-0.

50. Fernandes, A. P. et al. Expression profiles of thioredoxin family proteins in human lung cancer tissue: Correlation with proliferation and differentiation. Histopathology 55, 313–320 (2009).

51. Ran, D.-M., Zhang, Q.-W., Su, H.-L., Wang, C. & Gao, F.-H. Original Article Expression of thioredoxin reductase-1 and its effect in non-small cell lung cancer. Int J Clin Exp Med vol. 9 (2016).

52. Li, Z. et al. NQO1 protein expression predicts poor prognosis of non-small cell lung cancers. BMC Cancer 15, 207 (2015).

53. Li, Y. et al. Axl as a potential therapeutic target in cancer: Role of Axl in tumor growth, metastasis and angiogenesis. Oncogene (2009) doi:10.1038/onc.2009.212.

54. Jung, C. Y., Kim, S. Y. & Lee, C. Carvacrol targets AXL to inhibit cell proliferation and migration in non-small cell lung cancer cells. Anticancer Res. (2018) doi:10.21873/anticanres.12219.

55. HEPACAM inhibited the growth and migration of cancer cells in the progression of non-small cell lung cancer. - Abstract - Europe PMC. http://europepmc.org/article/med/26392113.

56. Lennon, F. E. et al. Overexpression of the μ-opioid receptor in human non-small cell lung cancer promotes akt and mTOR activation, tumor growth, and metastasis. Anesthesiology (2012) doi:10.1097/ALN.0b013e31824babe2.

57. Lennon, F. E. et al. The Mu opioid receptor promotes opioid and growth factor-induced proliferation, migration and Epithelial Mesenchymal Transition (EMT) in human lung cancer. PLoS One (2014) doi:10.1371/journal.pone.0091577.

58. Huang, C. Y. et al. CCL5 increases lung cancer migration via PI3K, Akt and NF-κB pathways. Biochem. Pharmacol. (2009) doi:10.1016/j.bcp.2008.11.014.

59. Sheng, H., Li, X. & Xu, Y. Knockdown of FOXP1 promotes the development of lung adenocarcinoma. Cancer Biol. Ther. (2019) doi:10.1080/15384047.2018.1537999.

60. Jin, D. et al. UBE2C, directly targeted by miR-548e-5p, increases the cellular growth and invasive abilities of cancer cells interacting with the EMT marker protein zinc finger e-box binding homeobox 1/2 in NSCLC. Theranostics 9, 2036–2055 (2019).

61. Li, H. et al. PTTG1 promotes migration and invasion of human non-small cell lung cancer cells and is modulated by miR-186. Carcinogenesis (2013) doi:10.1093/carcin/bgt158.

62. Xu, X. et al. Network analysis of DEGs and verification experiments reveal the notable roles of PTTG1 and MMP9 in lung cancer. Oncol. Lett. (2018) doi:10.3892/ol.2017.7329.

63. Wu, L. & Yang, L. The function and mechanism of HMGB1 in lung cancer and its potential therapeutic implications. Oncology Letters vol. 15 6799–6805 (2018).

64. Cao, J. et al. Prdx1 inhibits tumorigenesis via regulating PTEN/AKT activity. EMBO J. 28, 1505–1517 (2009).

65. Zhang, X. Q., Chen, G. P., Wu, T., Yan, J. P. & Zhou, J. Y. Expression and clinical significance of Ezrin in non-small-cell lung cancer. Clin. Lung Cancer 13, 196–204 (2012).

66. Saygideğer-Kont, Y. et al. Ezrin Enhances EGFR Signaling and Modulates Erlotinib Sensitivity in Non-Small Cell Lung Cancer Cells. Neoplasia (2016) doi:10.1016/j.neo.2016.01.002.

67. Fan, X. et al. B-Myb mediates proliferation and migration of non-small-cell lung cancer via suppressing IGFBP3. Int. J. Mol. Sci. (2018) doi:10.3390/ijms19051479.

68. Jin, Y. et al. B-Myb is up-regulated and promotes cell growth and motility in non-small cell lung cancer. Int. J. Mol. Sci. (2017) doi:10.3390/ijms18060860.

69. Yu, Y. H. et al. Network biology of tumor stem-like cells identified a regulatory role of cbx5 in lung cancer. Sci. Rep. (2012) doi:10.1038/srep00584.

70. Shao, C. et al. Essential role of aldehyde dehydrogenase 1A3 for the maintenance of non-small cell lung cancer stem cells is associated with the STAT3 pathway. Clin. Cancer Res. (2014) doi:10.1158/1078-0432.CCR-13-3292.

71. Hung, M. H. et al. SET antagonist enhances the chemosensitivity of non-small cell lung cancer cells by reactivating protein phosphatase 2A. Oncotarget (2016) doi:10.18632/ONCOTARGET.6313.

72. Katono, K. et al. Prognostic significance of MYH9 expression in resected non-small cell lung cancer. PLoS One (2015) doi:10.1371/journal.pone.0121460.

73. Leung, E. L. H. et al. Non-small cell lung cancer cells expressing CD44 are enriched for stem cell-like properties. PLoS One (2010) doi:10.1371/journal.pone.0014062.

74. Fu, H. et al. Aldolase A promotes proliferation and G 1 /S transition via the EGFR/MAPK pathway in non-small cell lung cancer. Cancer Commun. (London, England) (2018) doi:10.1186/s40880-018-0290-3.

75. Chang, Y. C. et al. Feedback regulation of ALDOA activates the HIF-1α/MMP9 axis to promote lung cancer progression. Cancer Lett. (2017) doi:10.1016/j.canlet.2017.06.001.

76. Tang, F. et al. SBI0206965, a novel inhibitor of Ulk1, suppresses non-small cell lung cancer cell growth by modulating both autophagy and apoptosis pathways. Oncol. Rep. (2017) doi:10.3892/or.2017.5635.

77. Yu, H. et al. SET domain containing protein 5 (SETD5) enhances tumor cell invasion and is associated with a poor prognosis in non-small cell lung cancer patients. BMC Cancer (2019) doi:10.1186/s12885-019-5944-2.

78. Wang, H. et al. Overexpression of ELF3 facilitates cell growth and metastasis through PI3K/Akt and ERK signaling pathways in non-small cell lung cancer. Int. J. Biochem. Cell Biol. (2018) doi:10.1016/j.biocel.2017.12.002.

79. Jian, H., Zhao, Y., Liu, B. & Lu, S. SEMA4b inhibits MMP9 to prevent metastasis of non-small cell lung cancer. Tumor Biol. (2014) doi:10.1007/s13277-014-2409-8.

80. Jian, H., Zhao, Y., Liu, B. & Lu, S. SEMA4B inhibits growth of non-small cell lung cancer in vitro and in vivo. Cell. Signal. (2015) doi:10.1016/j.cellsig.2015.02.027.

81. C, M. et al. Protease Serine S1 Family Member 8 (PRSS8) Inhibits Tumor Growth In Vitro and In Vivo in Human Non-Small Cell Lung Cancer. Oncol. Res. 25, 781–787 (2016).

82. Liu, Y. et al. The transcription factor DEC1 (BHLHE40/STRA13/SHARP-2) is negatively associated with TNM stage in non-small-cell lung cancer and inhibits the proliferation through cyclin D1 in A549 and BE1 cells. Tumor Biol. (2013) doi:10.1007/s13277-013-0697-z.

83. Xia, P., Wang, W. & Bai, Y. Claudin-7 suppresses the cytotoxicity of TRAIL-expressing mesenchymal stem cells in H460 human non-small cell lung cancer cells. Apoptosis (2014) doi:10.1007/s10495-013-0938-z.

84. Lu, Z. et al. Claudin-7 inhibits human lung cancer cell migration and invasion through ERK/MAPK signaling pathway. Exp. Cell Res. 317, 1935–1946 (2011).

85. Song, S. et al. Gene silencing associated with SWI/SNF complex loss during NSCLC development. Mol. Cancer Res. (2014) doi:10.1158/1541-7786.MCR-13-0427.

86. Orvis, T. et al. BRG1/SMARCA4 inactivation promotes non-small cell lung cancer aggressiveness by altering chromatin organization. Cancer Res. (2014) doi:10.1158/0008-5472.CAN-14-0061.

87. Gastonguay, A. et al. The role of Rac1 in the regulation of NF-kB activity, cell proliferation, and cell migration in non-small cell lung carcinoma. Cancer Biol. Ther. (2012) doi:10.4161/cbt.20082.

88. Zhou, Y. et al. Rac1 overexpression is correlated with epithelial mesenchymal transition and predicts poor prognosis in non-small cell lung cancer. J. Cancer 7, 2100–2109 (2016).

89. Guo, Y. et al. Cyclophilin A promotes non-small cell lung cancer metastasis via p38 MAPK. Thorac. Cancer 9, 120–128 (2018).

90. Ying, Q. L. et al. The ground state of embryonic stem cell self-renewal. Nature (2008) doi:10.1038/nature06968.

91. Schlesinger, S. & Meshorer, E. Open Chromatin, Epigenetic Plasticity, and Nuclear Organization in Pluripotency. Developmental Cell (2019) doi:10.1016/j.devcel.2019.01.003.

92. Ying, Q. L. & Smith, A. The Art of Capturing Pluripotency: Creating the Right Culture. Stem Cell Reports (2017) doi:10.1016/j.stemcr.2017.05.020.

93. Silva, J. et al. Nanog Is the Gateway to the Pluripotent Ground State. Cell (2009) doi:10.1016/j.cell.2009.07.039.

94. Festuccia, N. et al. Esrrb is a direct Nanog target gene that can substitute for Nanog function in pluripotent cells. Cell Stem Cell (2012) doi:10.1016/j.stem.2012.08.002.

95. Fukushima, A. et al. Characterization of functional domains of an embryonic stem cell coactivator UTF1 which are conserved and essential for potentiation of ATF-2 activity. J. Biol. Chem. (1998) doi:10.1074/jbc.273.40.25840.

96. Carey, B. W., Markoulaki, S., Beard, C., Hanna, J. & Jaenisch, R. Single-gene transgenic mouse strains for reprogramming adult somatic cells. Nat. Methods (2010) doi:10.1038/NMETH.1410.

97. Cheung, H.-H., Liu, X. & Rennert, O. M. Apoptosis: Reprogramming and the Fate of Mature Cells. ISRN Cell Biol. (2012) doi:10.5402/2012/685852.

98. Tirosh, I. et al. Single-cell RNA-seq supports a developmental hierarchy in human oligodendroglioma. Nature 539, 309 (2016).

99. Cao, J. et al. Comprehensive single-cell transcriptional profiling of a multicellular organism. Science (80-.). (2017) doi:10.1126/science.aam8940.

100. Baron, M. et al. A Single-Cell Transcriptomic Map of the Human and Mouse Pancreas Reveals Inter- and Intra-cell Population Structure. Cell Syst. (2016) doi:10.1016/j.cels.2016.08.011.

101. Kowalczyk, M. S. et al. Single-cell RNA-seq reveals changes in cell cycle and differentiation programs upon aging of hematopoietic stem cells. Genome Res. (2015) doi:10.1101/gr.192237.115.

102. Geirsdottir, L. et al. Cross-Species Single-Cell Analysis Reveals Divergence of the Primate Microglia Program. Cell (2019) doi:10.1016/j.cell.2019.11.010.

103. Navin, N. E. The first five years of single-cell cancer genomics and beyond. Genome Res. 25, 1499–1507 (2015).

104. Qiu, P. Embracing the dropouts in single-cell RNA-seq analysis. Nat. Commun. 11, 1–9 (2020).

105. Jordan, C. T., Guzman, M. L. & Noble, M. Cancer Stem Cells. N. Engl. J. Med. 355, 1253–1261 (2006).

106. Wagner, D. E. et al. Single-cell mapping of gene expression landscapes and lineage in the zebrafish embryo. Science (80-.). 360, 981–987 (2018).

107. Mootha, V. K. et al. PGC-1α-responsive genes involved in oxidative phosphorylation are coordinately downregulated in human diabetes. Nat. Genet. (2003) doi:10.1038/ng1180.

108. Li, B. & Dewey, C. N. RSEM: accurate transcript quantification from RNA-Seq data with or without a reference genome. BMC Bioinformatics 12, 323 (2011).

109. Langmead, B., Trapnell, C., Pop, M. & Salzberg, S. L. Ultrafast and memory-efficient alignment of short DNA sequences to the human genome. Genome Biol. 10, (2009).

